# Genetic Modulation of Lifespan: Dynamic Effects, Sex Differences, and Body Weight Trade-offs

**DOI:** 10.1101/2025.04.27.649857

**Authors:** Danny Arends, David G. Ashbrook, Suheeta Roy, Lu Lu, Zachary Sloan, Arthur G. Centeno, Kurt H. Lamour, João Pedro de Magalhães, Pjotr Prins, Karl W. Broman, Saunak Sen, Sarah J. Mitchell, Michael R. MacArthur, Özlem Altintas Akin, Johan Auwerx, Amandeep Bajwa, Vivian Diaz, David E. Harrison, Randy Strong, James F. Nelson, Khyobeni Mozhui, Evan G. Williams, Richard A. Miller, Robert W. Williams

## Abstract

The dynamics of lifespan are shaped by DNA variants that exert effects at different ages. We have mapped genetic loci that modulate age-specific mortality using an actuarial approach. We started with an initial population of 6,438 pubescent siblings and ended with a survivorship of 559 mice that lived to at least 1100 days. Twenty-nine *Vita* loci dynamically modulate the mean lifespan of survivorships with strong age- and sex-specific effects. Fourteen have relatively steady effects on mortality while other loci act forcefully only early or late in life and with polarities of effects that invert. A distinct set of 19 *Soma* loci shape the *negative* correlation between weights of young adults with their life expectancies—much more strongly so in males than females. Another set of 11 *Soma* loci shape the *positive* correlation between weights at older ages with life expectancies. The *Vita* and *Soma* loci share 289 age-dependent epistatic interactions (LODs ≥3.8) but fewer than 4% are common to both sexes. We provide two examples of how to move from maps toward potential mechanisms. Our findings provide an empirical bridge between evolutionary theories on aging and genetic and molecular causes. These loci and their interactions are key to begin to understand the impact of interventions that may extend healthy lifespan in mice and even in humans.

---

We do not yet understand genetic, molecular, cellular, and organismal processes that shape intrinsic variability in rates of aging, lifespan, and all-cause mortality in humans, mice, or other model organisms such as *Drosophila*^1–19^. While thousands of variants modulate risks of age-related diseases^20,21^, the great majority influence proximate causes of death but not the core mechanisms that influence aging rates *per se*^22–26^. To disentangle causes from consequences and to define variants that modulate mortality across entire lifespans, we developed an actuarial approach that maps variation in mean lifespan of progressively older survivorships; from those that include all mice from pubescence upward, to those made up only of mice that lived to at least 1100 days. This is equivalent to a range of survivorships starting from 12 to 94 years in humans^27,28^. This approach has been tested before^5,6,29–33^, but has not been applied systematically in any organism. The closest human parallels are biometric studies of age-dependent changes in heritability of lifespan in Scandinavian twin cohorts born between 1870 and 1910^34,35^. Our study is a complementary dissection but at the level of discrete genetic effects—59 well-defined loci that dynamically interact to influence mortality rates and lifespan. We address four sets of questions in biodemographics and geroscience:

1. What are the chromosomal coordinates of genetic variants that influence mortality rates, and how and when do they act? Which have effects consistent with modulating aging rates, which have transient effects consistent with specific diseases, which have effects only late in life?
2. How do genetic variants align with sex differences in mortality rate during and after the reproductive phase of life? Is there evidence of genotype-by-sex (G×S) and genotype-by-genotype (G×G) interactions or even antagonism?
3. What loci account for the strong negative association between larger body size early in life and shortened lifespan? Are these loci an independent set?
4. How do the dynamics of action of loci and their epistatic interactions support or refute predictions made by major evolutionary and life history theories of aging—the *mutation accumulation theory*^36,37^*, the antagonistic pleiotropy* theory^38^, and the *longevity-assurance*/*disposable soma theory*^17,39^?

To dissect the genetic architecture of lifespan we relied on the largest study of mouse aging—the National Institute on Aging’s *Interventions Testing Program* (ITP)^40^. For over two decades, teams at three sites have used a highly diverse population of full genetic siblings called the *University of Michigan Heterogeneous Cohort 3* (UM-HET3) to study effects of drug interventions on longevity^40–42^. These UM-HET3 mice are created by crossing hybrids of four fully inbred progenitors—BALB/cByJ (cBy or *C*), C57BL/6J (B6 or *B*), C3H/HeJ (C3 or *H*), and DBA/2J (D2 or *D*)—that have a longevity range from 600 days in D2 to 900 days in B6^43,44^. UM-HET3 progeny segregate for ∼11 million DNA variants—a level of heterogeneity comparable to that of humanity^45^. Between 2004 and 2013, ITP teams took tissues from 13,200 pubescent mice. We have used 6,438 of these samples for genetic analysis—excluding only those mice treated with dietary supplements such as rapamycin that significantly extend lifespan^41^. All siblings were born and aged to natural death in three pathogen-free vivaria at the Jackson Laboratory in Bar Harbor, Maine, the University of Michigan in Ann Arbor, Michigan, and the University of Texas Health San Antonio, Texas. Our findings are robust with respect to a decade-long experiment and marked environmental differences among these three sites.

In earlier work we mapped lifespan using 3,055 mice and detected seven lifespan loci^46^. Here we have doubled sample size by adding 3,383 mice given supplements that did *not* affect lifespan. We quadrupled numbers of genetic markers and introduced a new actuarial method. The result has been a four-fold increase in numbers of lifespan loci *(Vita* type). In addition, we have mapped 30 loci of a new type that specifically influence correlations between body weight and life expectancies in males and females (*Soma* type). Finally, our sample size has enabled us to comprehensively analyze epistatic interactions and their sex and age-dependent effects among all loci.

The dynamics of *Vita* and *Soma* loci are highly complex. Some act in early or late survivorships. A subset modulate expectancies more uniformly and include DNA variants that may act as pacemakers of aging^47^. Many have striking and even counterbalanced sex differences, as do almost all epistatic partnerships. Some have effects predicted by the theory of antagonistic pleiotropy^36,48,49^, while others have effects long after the reproductive phase and are consonant to different degrees with both the *mutation accumulation* theory and the longevity-assurance/*disposable soma* theory^2,17,38,39,50^. With persistence, many of these loci should be reducible to well defined molecular and cellular mechanisms.

## Results

We analyzed life expectancies of 6,438 mice belonging to 72 nested survivorships each of which was generated by truncating upward in 15-day steps. The base survivorship in Fig. 1a includes all mice that entered the study between 2004 and 2013 at puberty. The last survivorship includes only a subset of 559 of these mice (8.7%) with lifespans between 1100 and 1456 days (Fig. 1a,b). We refer to each survivorship by its truncation age (T-age). At the base truncation of T-42 days (T_42_) the sex difference in mean lifespans of UM-HET3 mice is 81 days (Fig. 1b); that for males is 806 ± 210 SD days; that for females is 887 ± 175 SD days. This difference is stable from T_42_ to T_215_ because so few mice die in the first 215 days of life—20 males and 13 females. In contrast, between 215 and 410 days many more males die than do females—204 versus 18. This striking sex difference in early mortality produces the symmetric inflections in the actuarial effect size plot in Fig 1b. They are symmetric because values are normalized to the mean for every T-age. By T_740_, lifespans of male and female survivorships have converged precisely—946 ± 2.7 and 946 ± 2.4 SE days (Fig. 1b). The terminal survivorship of 277 males and 282 females have lifespans of 1166 ± 3.2 and 1159 ± 3.3 (s.e.) days.

**Fig 1.**
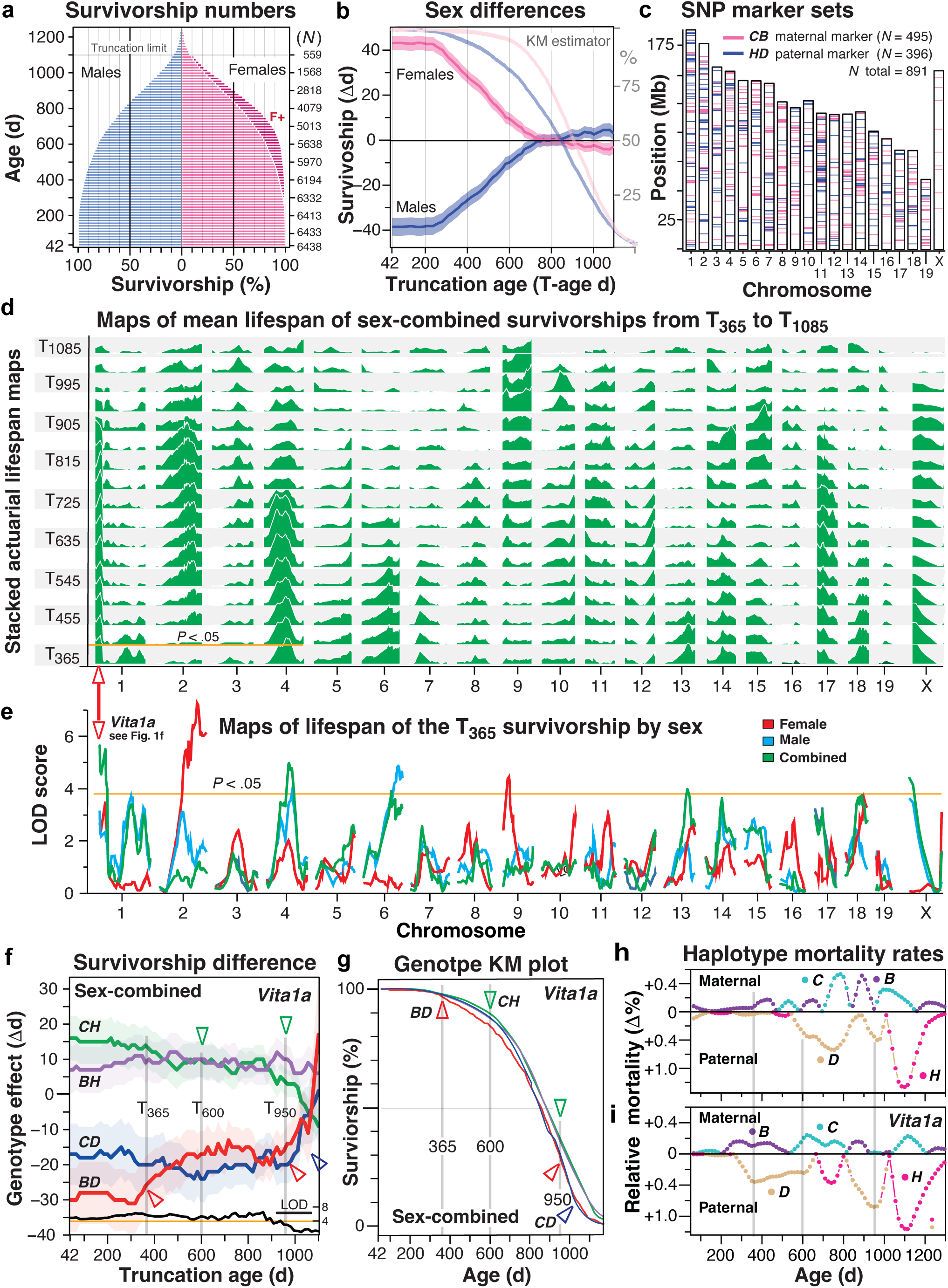
Mortality, sex differences, and *Vita* survivorship loci. **a,** Survivorships of mice stratified by the minimum inclusion age starting at a population of 6,438 at our base truncation age (T-age) of T_42_ and extending to a high of T_1100_, This final survivorship has 559 mice. The first death was at 46 days, the last death at 1456 days. 364 more males (*blue bars*) were entered into the study, but by T_530_ n/sex was matched. *F+* denotes the proportional survival advantage of females (darker red bars, 4% at T_365_). Truncation ages (left *y-axis*) and numbers of survivors on the right *y*-axis. **b,** Sex differences in mean lifespans of the T_42_ to T_1100_ survivorships. The 81-day difference at T_42_ is neutralized by T_725_. Kaplan-Meier (KM) estimators for both sexes are superimposed. **c,** Ideogram of chromosomes and approximate megabase (Mb, GRCm38/mm10) of SNPs used to define maternal (*C* vs *B*) and paternal haplotypes (*H* vs *D*). **d,** Stacked survivorship maps for sex-combined data from T_365_ to T_1085_. We provide corresponding versions of panel **d** stratified by sex and over a wider age range in Extended Data 1. **e,** The map of T_365_ corresponds to the lowest tier of panel **d** but now with resolution of female (*red*), male (*blue*), and combined (*green*) plots. The *red triangle* marks the *Vita1a* locus. **f**, Actuarial genetic effect size plots from T_42_ to T_1100_. The *y*-axis defines the mean lifespan differences of survivorships for the four genotypes as a function of truncation age (*x-*axis) relative to the average of all four genotypes. LOD scores are plotted above the *x*-axis with an orange line at *P* <.05. The trios of faint vertical lines in **f-h** are for help in comparing these three different plot types. **Red** and **blue triangles** mark inflection points in mean lifespan of *BD* and *CD* survivorship at T_365_ and T_950_. **g,** KM plot of survival stratified by the four genotypes with arrowheads. **h,i,** Plots of age-dependent differences in relative mortality rates at *Vita1a* in females (**h**) and males (**i**). Each panel is divided into an upper maternal block with mortality rates of the *C* and *D* maternal haplotypes, and a lower paternal block with relative mortality rates of paternal *D* and *H* haplotypes. Deviations away from zero (no difference) in either direction signify a relative increase in mortality for that haplotype relative to the alternative haplotype. Note that the *D* haplotype is strongly disadvantageous only before 900 days, whereas the *H* haplotype is disadvantageous after 1000 days. We used a LOESS with α of 0.2 over 75-day mean haplotype mortality counts. Gray lines for comparison across sexes and with **f** and **g**.

### 29 *Vita* loci defined by actuarial dissection of 72 survivorships

We resolved loci modulating mean lifespans of survivorships at a genome-wide false discovery rate <0.05 after applying a stringent Bonferroni correction and a Cauchy correction for actuarial time-series analysis (Table 1, Fig. 1, Methods). The mapping was stratified by sexes and in combination (Fig. 1d,e, Extended Data 1). Individual loci have average effects of 36 ± 12 days (d) on life expectancies that depend on sex and T-age (Table 1, Fig. 1c-f, Extended Data 1–3). Peak effects explain 2.5 ± 0.9% of variance in females and 3.2 ± 0.5% in males. Linkage ranges from a low of 4.1 LOD (*Vita5a*) to a high of 9.0 in males (*Vita4a*) and to a high of 8.1 in females (*Vita2c*). Confidence intervals average 35 ± 19 Mb SD^51^ with the smallest two under 10 Mb (Table 1, Fig. 1d-f). *Vita* loci have age-delimited actuarial effects that average 349 ± 229 days SD (Table 1, average of *Duration* column) and durations that range from 45 to 900 days.

**Table 1.**
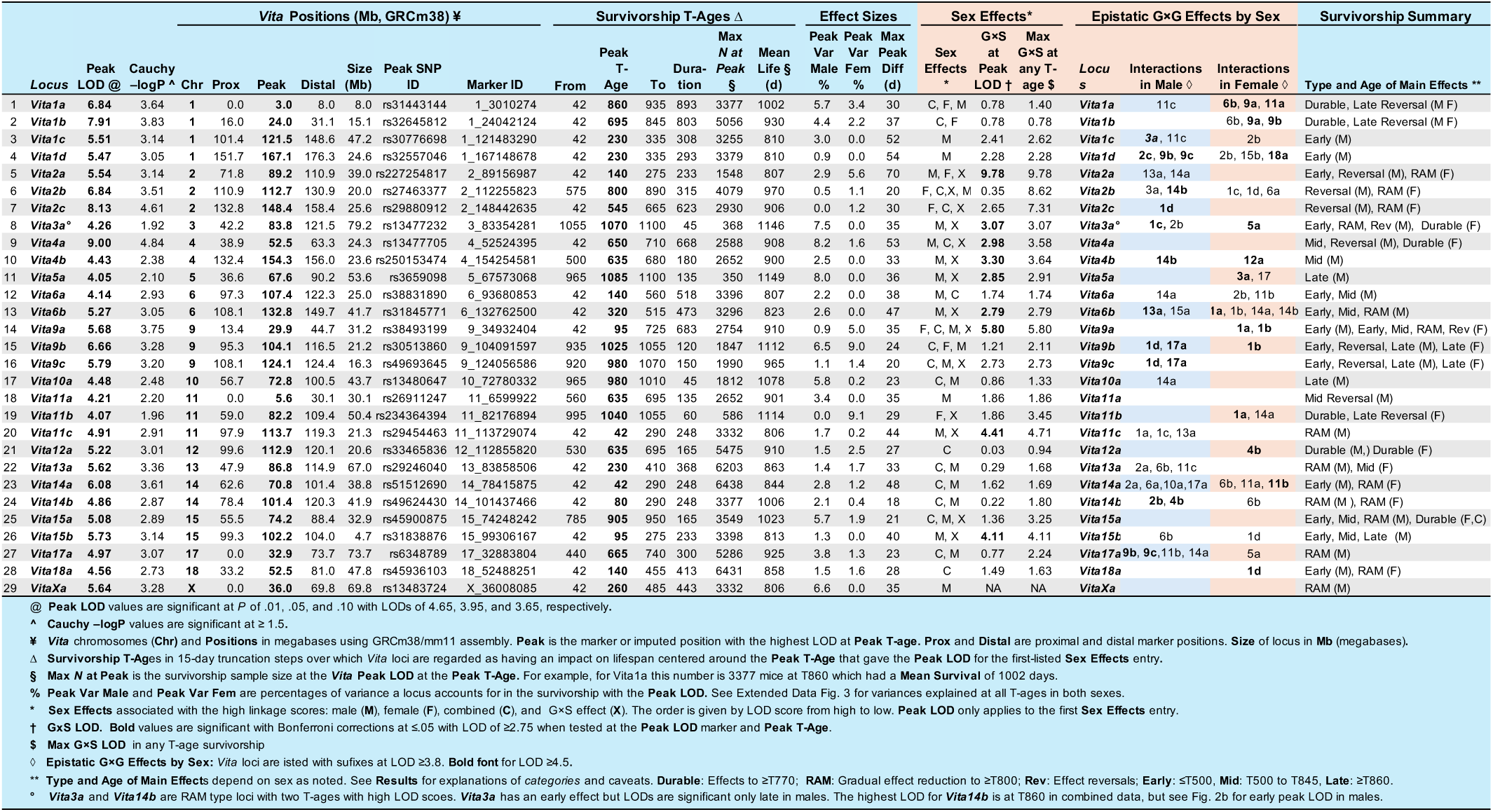
29 *Vita* loci modulate age-dependent mortality rates and life expectancies.

### Dynamics of *Vita* loci

Effects of *Vita* loci on lifespan are highly age-dependent (Fig. 1; Table 1, right-most column; Extended Data 1–4). *Vita1a* and *Vita1b* have relatively stable actuarial effects from T_42_ to T_890_ (Figs 1f, 2a). In contrast, *Vita14b* has a gradual reduction in its effects across survivorships (Fig. 2b) with converging slopes of both *BD* and *CH* genotypes from T_230_ to T_1100._ Other loci have transient effects. *Vita1c* acts only in survivorships that include adults younger than 500 days (Fig. 2c) whereas *Vita9c* acts only in the oldest survivorships (Fig. 2d). *Vita4a* has effects that reverse between T_410_ and T_800_ in males (Fig. 2e,f). Inflection points in these actuarial plots correspond to ages on Kaplan-Meier estimators (Fig. 1g) at which genotype classes diverge or converge—at which rates of mortality change (Extended Data 2–5). Figs. 1h and 2f highlight strong age-dependent differences in mortality rates of maternal and paternal haplotypes on risk of dying by sex and age in which the *D* haplotype contributes to higher mortality earlier in life (for similar data for all *Vita* loci see Extended Data 4). This explains the actuarial benefit noted even in the earliest survivorships of inheriting an *H* haplotype (Fig. 1f). An important conclusion: the biological impact of DNA variants in *Vita* loci are generally greatest at inflections in actuarial plots and at peaks and troughs in mortality rate plots (Fig. 2a-f, Extended Data 2, 3, 4). These are T-age ranges over which lifespans of survivorships swing by up to 15 days over just 45 days (Fig. 2e). In contrast, those *Vita* loci that have a comparatively constant slope from T_42_ to beyond T_905_ (Fig. 2b) behave as expected of rate of aging modulators^10,47^.

**Fig 2.**
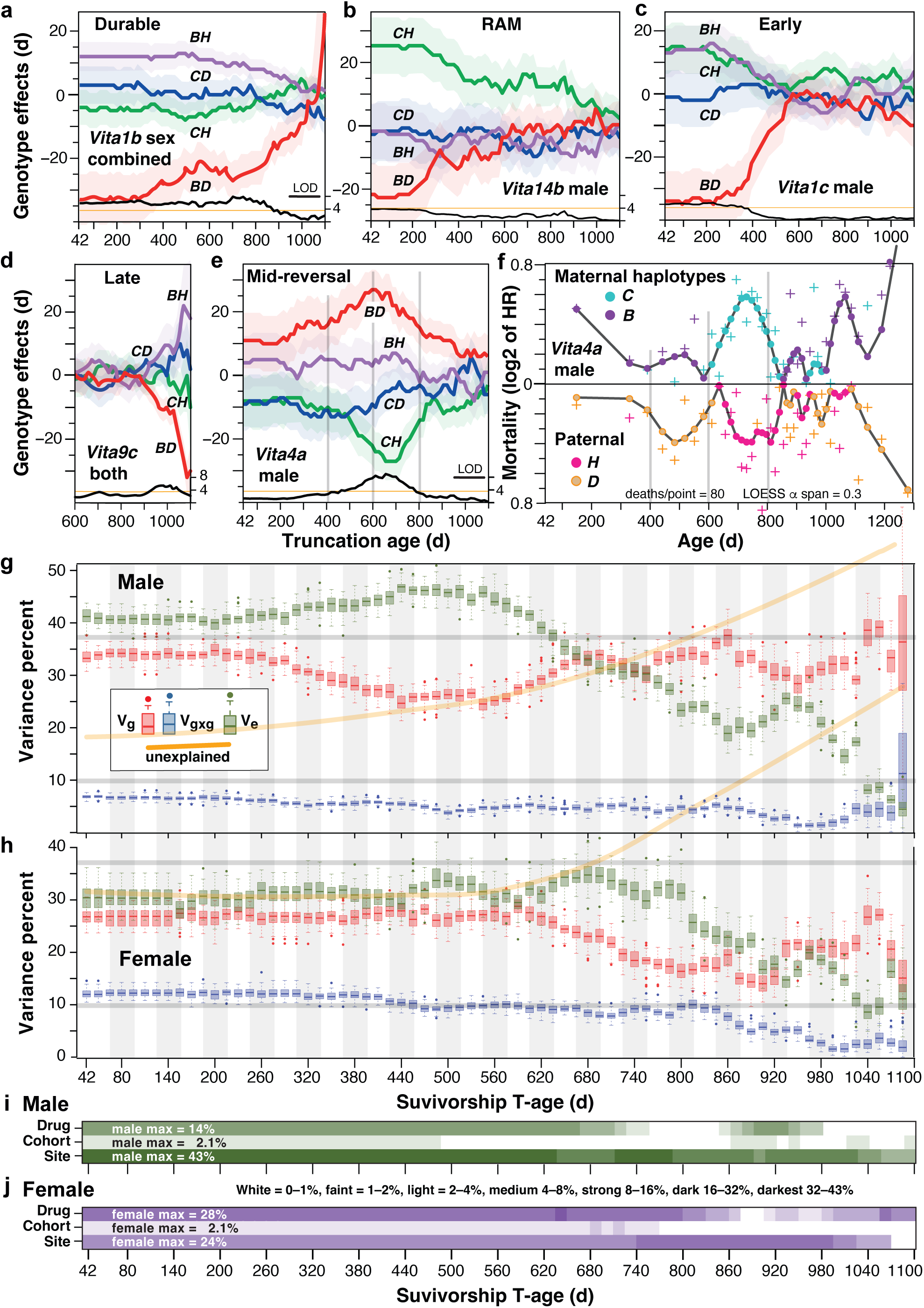
Dynamics of *Vita* loci, their contributions to heritability, and source of variance. **a–f,** Genetic effect plots define differences in the mean lifespan of survivorships for the four genotypes (*BH*, *BD*, *CH*, *CD*) for different categories of *Vita* loci. The effects of genotypes in days are marked by colored lines on the *y*-axis (relative difference from the average). Two-letter labels define genotypes (pairs of haplotypes). LOD scores are plotted along the bottom of each plot as a black trace above the T-age *x*-axis. *Vita* locus names are marked in all panels. Extended Data 2 provides these plots for sex-combined data, females, and males. **a,** *Vita1b* has durable and relatively uniform actuarial effects that extend from T_42_ to T_900_ survivorships. The initially negative effect of the *BD* genotype in most survivorships is caused by higher mortality visible starting in the T_855_ survivorship (*Results* for explanation). **b,** Vita*14b* is a candidate rate of aging modulator that has its highest LOD scores in the base male survivorship at T_42_ that reflects a life-long and uniform contrast in mortality rates between *BD* and *CH* genotypes. All modes of display are compared for this locus in Extended Data 5, enabling readers to comparing survivorship plots with age-dependent mortality. **c**, *Vita1c* has early effects in male mortality from the infections at T_365_ and T_590_. **d**, *Vita9c* is an example of a locus with late effects after T_935_ due to complex sex difference (Extended Data 2q). **e** *Vita4a* has a striking reversal of genotype effects between T_450_ and T_810_—marked by three vertical gray lines in both panels—that is caused by offset waves of mortality in males: an early phase of *C* haplotype mortality starting at 400 days (**f**) and a delayed phase of *H* haplotype mortality starting at 560–600 days that both peak at roughly 750 to 800 days. **f,** Age-dependent hazard ratios for males at *Vita4a.* Each cross-hair point represents 80 mortality events over an age range centered on the point that are either *C* or *B* maternal haplotypes (top) or H or D paternal haplotypes. Points were smoothed using a LOESS with a span of 0.3 (Methods). **g,h,** Comparison of the variance explained across all survivorships specifically by peak markers. The pink boxplots (*Vg*) are estimates of variance that can be explained by all 29 *Vita* loci. The blue boxplots are similar estimates of variance explained by the subset of *Vita*-*Vita* epistatic interactions (*Vgxg*) defined in a larger bold font in Fig. 5b. The green boxplots (*Ve*) estimate the summed environmental and experimental variance further broken down in panels **i** and **j**. The orange lines are smoothed averages of the variance that we cannot explain, including gene-by-environmental effects and higher-order genetic factors such as indirect genetics effects of co-housed mice. **h,** Female components as above, but note the significantly lower variance values compared to males. The gray horizontal lines at 10% and 37.5% in (**g)** and (**h)** are to help compare variance levels between sexes. **i,j,** Breakdowns of non-genetic sources of variance. Rows labeled *Drug* are variance estimates of dietary supplements versus the control diet. *Cohort* is variance associated with year of the production of a cohort—nine years total. *Site* is variance attributable to differences among the three vivaria. For a display of *Vg* of individual *Vita* loci for all survivorships see Extended Data 6.

We have categorized the dynamics of *Vita* loci into four categories (Table 1, far right side) using all of the data and plots provided in Extended Data 2–4.

**1. Loci with durable effects**. Five loci have relatively constant actuarial effects on mean life expectancies of survivorships from as early as the T_42_ base survivorship out to the T_750_ survivorship in one or both sexes (Table 1). *Vita1a* (Fig. 1f) is an excellent example. The actuarial durability and consistency of these loci means they are useful predictors of life expectancy even in adolescence.
**2. Loci with steadily diminishing effects**. These loci are characterized by uniform slopes in plots that start as early as 215 days in males and extend at least to the T_750_ survivorship in either sex. We list 14 of these loci as potential rate of aging modulators (RAMs) using a liberal definition of “uniform slopes”, but half would be accepted at a higher stringency of an age-independent hazard ratio for over 500 days. Their actuarial effects can converge toward zero difference in late survivorships. *Vita14b* (Fig. 2b) is a good example in which the initially high survivorship difference in males between *CH* and *BD* genotypes fades from T_305_ to T_1100_. This effect is due to high mortality of *B* and *D* haplotype carriers up to about 800 days followed by higher mortality of *C* and *H* carriers beyond that age (Extended Data 5). The T_42_ survivorship integrates lifelong mortality, whereas the later survivorships only integrate late phase mortality events driven by *C* and *H* haplotypes.
**3. Loci with age-range restricted effects.** This category includes loci with action limited to early, middle, or late survivorships (Fig. 2c-f). We define three subtypes.

**a. Early-age loci have strongest effects before T_500_**. The 15 loci of this type are just over three times more common in males than females (Table 1). The best examples are *Vita6a* (Fig. 2c), *Vita1c*, *Vita1d*, *Vita2a* (Extended Data 2c, 3c2), *Vita2b* (Fig. 3c-f), and *Vita18a* (Extended Data 2ab, 3ab). For example, the 50-day spread in lifespans of carriers of the *BH* and *BD* genotypes at *Vita1c* (Fig. 2c) is restricted to males (Extended Data. 2c, 3c1, 3c2). Exceptions are notable because polarities of genotype effects are flipped between sexes at four of these loci—*Vita2b*, *Vita9a*, *Vita11b*, and *Vita18a* (Fig. 3c-f, Extended Data 2n,s,ab, 2n1,s1, ab1). Only *Vita13*a has effect polarities shared by both sexes (Extended Data 2v, 3v1). The male bias in this type of locus is almost always linked to higher mortality over the range of T_200_ to T_700_ survivorships (Fig.1a,b, Extended Data 4).
**b. Mid-survivorship loci have strong effects from T_500_ to T_845_**. There are six loci of this type. Examples include *Vita12a* and *Vita2c* (Fig. 2c,d). *Vita4a* is an impressive example in which there is a strong biphasic fluctuation in the mortality rate of *BD* and *CH* genotype classes that is restricted to T_500_ to T_710_ male survivorships (Fig. 2e, Extended Data 2i, 3i1, 3i2). This odd actuarial effect is due to offset waves of mortality—the first *D* haplotype wave of deaths in males starting at 400 days (vertical line in Fig. 2f), followed at 600 days by waves of higher *C* and *H* haplotype hazard ratios. At 1000 days both *B* and *D* haplotypes have high relative mortalities and hazard ratios of nearly 2 (log2(HR) ∼ 1). The offsets in peaks of effects in the actuarial T-age plot (Fig. 2g) versus the mortality plot (Fig. 2f) is because the survivorships integrate mortality effects to the last death. The waves of genetic effects on mortality rates per se are much more obvious in the Fig 2f and for all other loci in Extended Data 4.
**c. Late-acting *Vita* loci have effects after T_860_**. Five loci have major effects after T_845_. *Vita9b* and *Vita9c* (Fig. 2d) have very late effects. *Vita5a* acts with progressively greater strength in male survivorships after T_730_ (Extended Data 2k). This category could include loci such as *Vita1a* that have strong late-life effects on mortality, but we do not include them for actuarial reasons—*Vita1a* predicts lifespan soon after birth, while this more formally defined late-acting category has no actuarial impact until the oldest survivorships.
**4. Loci with reversals of genetic effects.** Twelve loci have effects that reverse across survivorships. Reversals can involve a single genotype (the red *BD* trace in Fig. 2a) or two or more, for example *BD* and *CH* effects of *Vita4a* (Fig. 2e,f). These patterns are associated with age-dependent and highly variable hazard ratios.

**Fig 3.**
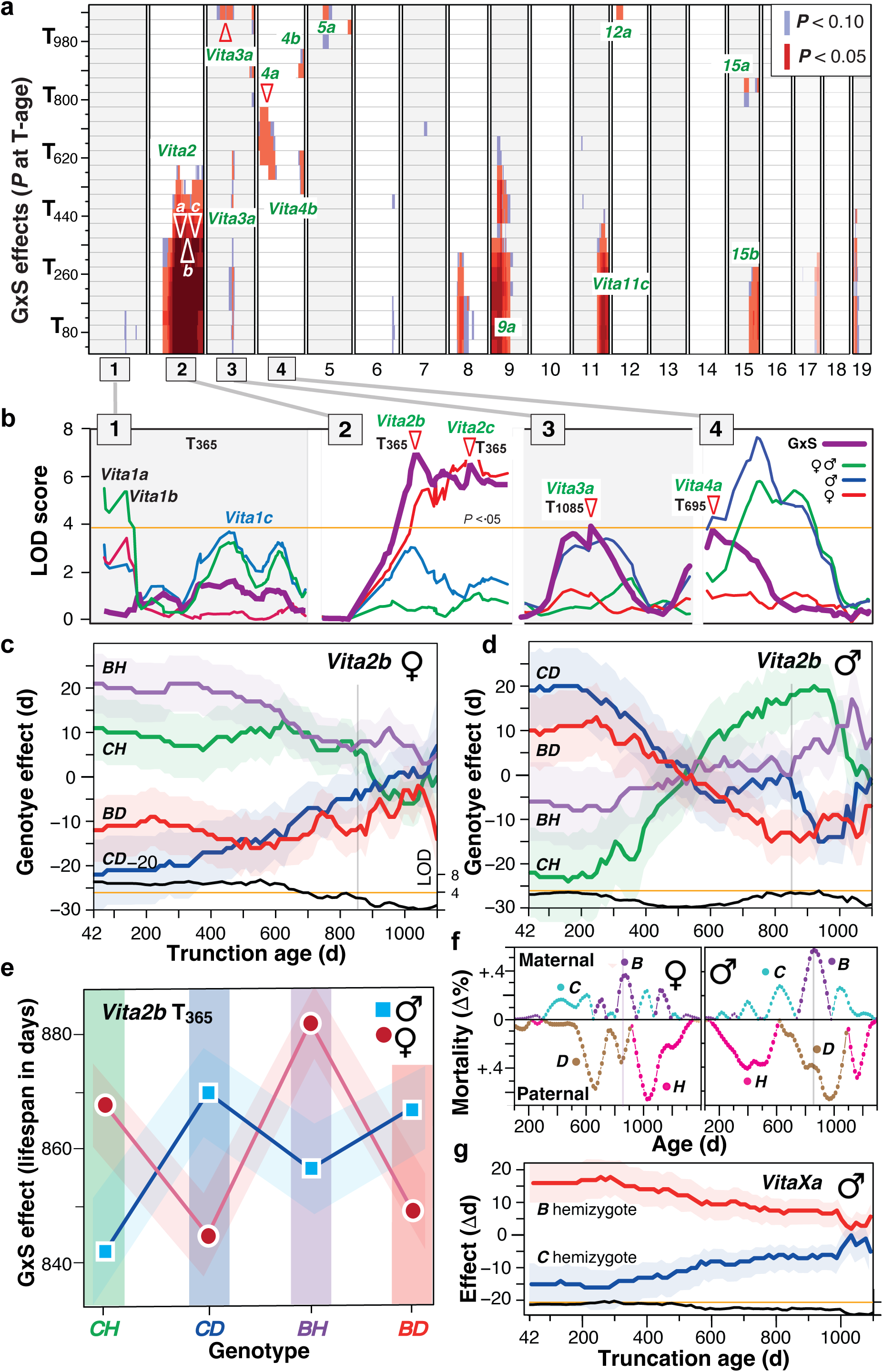
Antagonistic sex effects within *Vita* loci. **a,** Overview of all significant (red) and suggestive (light blue) locus-by-sex interaction effects (G×S) across all *Vita* loci in all survivorships. The G×S effect on Chr 8 is most likely a real interaction of a locus that has a main effect below our threshold. The G×S values in Table 1 are more discrete and only test top markers in the most significant survivorship. **b**, The first four chromosomes illustrate the range of sex differences in effects of *Vita* loci at T_365_. Red and blue lines signify female and male additive genetic effects, respectively (LOD scores), whereas the thicker purple lines signify the G×S effect, and the green lines are the “unsexed” or combined additive effect. Orange line is the genome-wide significance threshold at LOD 3.95. There are no G×S effects on Chr 1 (**b**, left). *Vita* loci with significant G×S effects on chromosomes 2, 3, and 4 are highlighted in green with the T-age of peak G×S at red triangles. **c,d**, *Vita2b* has strong G×S differences that define antagonistic (or complementary) effects on the two sexes. The notable reversals in life expectancies of genotypes in **d** are caused by higher mortality rates of carriers of the *C* and *H* haplotypes up to about 700 days, followed by higher mortality of *D* and *B* carriers from 800 to 1100 days (vertical lines at peak morality circa 850 days). **e**, The G×S difference and standard error of lifespans of males and females for the four genotypes. At T_365_ the C*H* and *BH* genotypes are advantageous for expectancies of females but the *CD* and *BD* haplotypes for males. **f**, Mortality rate differences (%) as a function of age, sex, and parental haplotypes as in Fig. 1h. In females, mortality rates of *D* carriers peak at 650 days, whereas *H* carriers do not peak until 1040 days. In males the *C* and *H* haplotypes have more uniform mortality distribution across lifespan whereas *B* and *D* haplotypes have a leptokurtic distributions that peak between 700 and 1050 days. **g,** The hemizygous *B* haplotype in males at *VitaXa* has a 30-day positive effect on lifespan compared to the *C* haplotype until T_300_.

**Fig 4.**
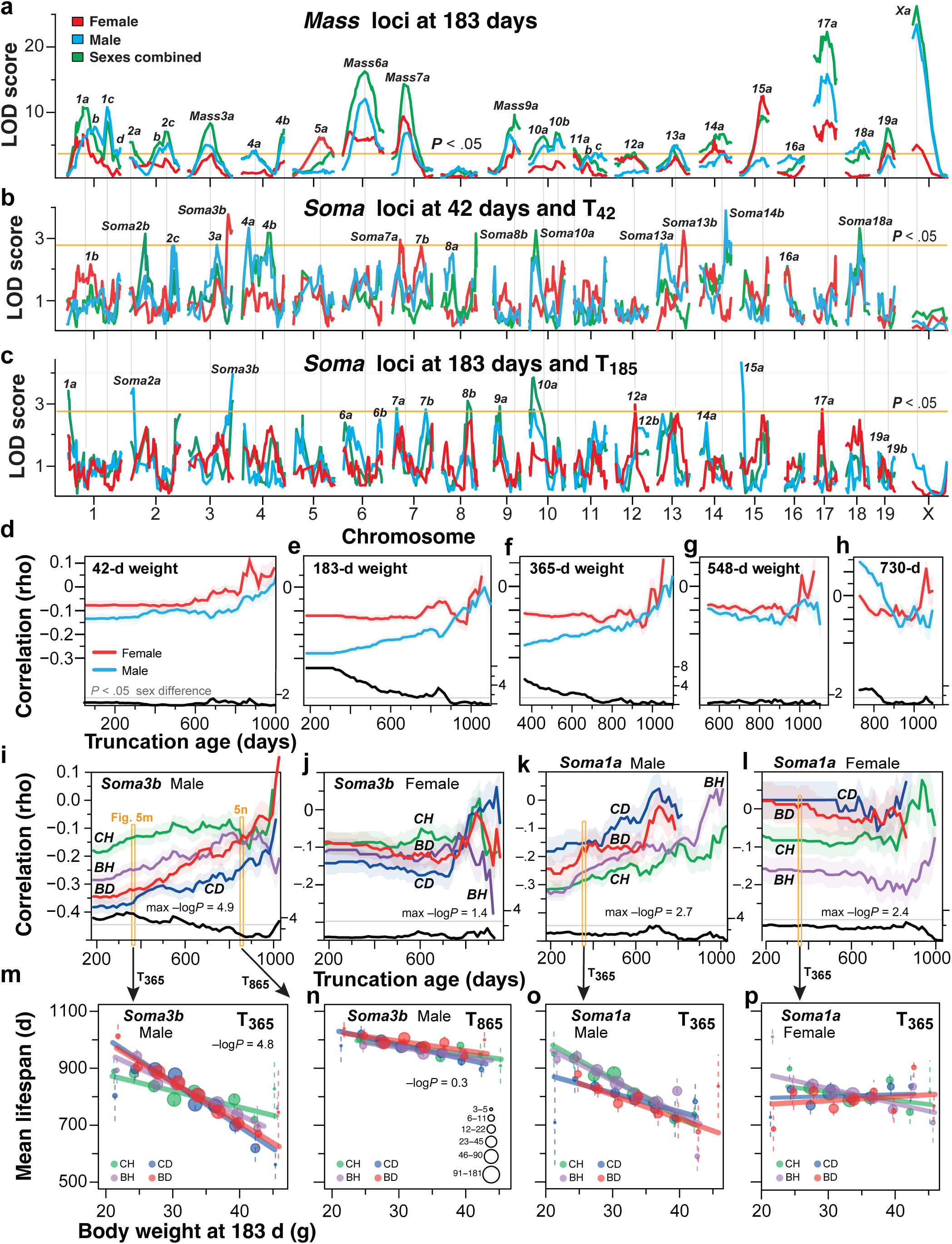
The genetic modulation of life expectancy by body weight is stronger in males than females. **a,** Conventional QTL maps of body mass (*Mass* loci) at 183 days for both sexes and combined. There are 25 significant *Mass* loci at T_183_ (see Extended Data 7 for all other ages). The yellow horizontal lines in **a**, **b** and **c** are genome-wide acceptance thresholds. The faint vertical gray lines at *Mass* peaks do not overlap significantly with *Soma* loci. **b,** Maps of the *Soma* loci that modulate correlations between body weight at 42 days and life expectancy in the T_42_ survivorship. Several loci are named despite being below threshold at this age or in (**c**). **c,** Corresponding *Soma* loci for body weight at 183 days (6 months) with life expectancy at T_185_. **d-h,** Actuarial plots for correlations of body weight at five ages (42, 183, 385, 558, and 730 days) for males and females. In **d** there is only a modest correlation of weight at 42 days with subsequent lifespan in any survivorship in either sex, but by 183 days in (**e**) there is a strong negative correlation in males and a significant sex difference (–log*P* of the sex difference is given above the *x*-axis). **g,** There is no sex difference at 548 days. **h,** Body weight at 730 days has a positive correlation in males but by the T_1040_ survivorship the female correlations are higher. **i,j,** Actuarial plots of correlations between weight at 183 days with subsequent survivorships for *Soma3b* for males **i** and females **j**. Correlations of the four color-coded genotypes differ significantly for males in the T_365_ survivorship (left orange bar) but not in the older T_665_ survivorship. The –logP *Soma* scores are given along the *x*-axis (black trace) with the maximum actuarial value. Note arrows from orange bars to panels **m** and **n**—cross-sectional views of these weight to life expectancy correlations at T_365_ that are highly significant in the T_365_ survivorship with a –logP of 4.8 but not significant in the T_865_ survivorship in **n**. **k,l,** Comparison of survivorship differences between males and females at *Soma1a*. *Soma1a* is close to the significance threshold in males in the T_710_ survivorship. Orange bars correspond to **o** and **p**. LOD scores (black lines) above the *x*-axis are significant at 2.75. Note the *y*-axis scale differences between sexes. **m-n,** Plots of *Soma3b* correlations (Spearman rank rho) in males in T_365_ and T_865_ survivorships. Survivorships were binned in 2-gram weight classes. The six circles sizes provide approximate sample size per class scaled to the log of the number. **o,p,** Comparison of male and female sex differences in *Soma* correlations that correspond to **k** and **l** at T_365_.

### Dynamics of heritability

The cumulative variance explained by all 29 *Vita* loci on lifespan is higher in males than females from T_42_ to T_320_ (Fig. 2g,h) but drops from its base at 40% to 30% by T_500_ (Fig. 2g, Extended Data 5). The decline is due to a wave of early male mortality. By T_650_ male heritability has rebounded and it continues to climb to 50% in the oldest survivorships. Female heritability is almost precisely 27% in all survivorships up to T_620_ but then drops to 20% by T_890_—450 days after the male low point. Like males, female heritability also recovers in the oldest survivorships, although still 10% lower than that of males. One important refinement on these sex differences is that the variance explained by epistatic interactions between *Vita* loci (Vgxg in Fig. 2g,h) is twice as high in females as it is in males—12% versus 6%, and this is true over the entire reproductive lifespan.

We estimated the impact of three non-genetic sources of variance: 1. treatment with or without a nominally ineffective dietary agent, 2. cohort’s year of birth, and 3. environmental differences at the three sites. Collectively, these variables differ significantly between sexes in magnitude and in their age-dependence (Fig. 2g,h). The overall level is ∼10% higher in males than females up to T_605_. In males the level peaks at 53% at T_440_ but drops in rough synchrony with the end of reproductive life in both male^52–54^ and female mice^55^. Less than 20% of variance in older survivorships is linked to these non-genetic sources—a counter-intuitive apparent reduction in sensitivity to exogenous factors later in life. The remaining unexplained sources of variance (orange lines in Fig. 2g,h) include uncontrolled environmental and technical factors and perhaps more importantly, higher order interaction effects that we have not modeled, including age-dependent changes in survivors per cage, and unmeasured indirect social genetic effects^56^.

Despite intense efforts to standardize protocols, variance that is attributable to *site* is up to 43% in males and 24% in females (Fig. 2i,j). In males, variance associated with receiving an ineffective drug versus no drug was much more modest than that of *site* (Fig. 2i) although in absolute terms these dietary supplements did increase lifespan at T_42_ by 40.2 ± 8.3 SE days. In contrast, in females the effect of these supplements was greater than that of site (Fig. 2j) and increased lifespan by 37 ± 6.7 SE days. This positive effect of the dietary supplement persisted to the T_680_ survivorship in males and to the T_800_ survivorship in females (Fig. 2i). Thereafter it turns negative at T_890_ in males (–11.7 ± 5.3 SE days, *P =* .03) and at T_1040_ in females (–10.2 ± 5.3 SE days, *P* = .06). Cohort year is not a significant source of variance in either sex.

### *Vita* loci have strong and complementary sex interaction effects

Fourteen *Vita* loci have strong gene-by-sex (G×S) interaction effects (Table 1, Fig. 3a). The complex of loci on Chr 2 is an extreme example (Fig. 3a-d). *CH* and *BH* genotypes at *Vita2b* confer an advantage to females that lasts to T_700_ (Fig. 3e), but that are disadvantageous to males through to T_365_. Polarities of the sex effects of genotypes are also reversed, with *CH* and *BH* genotypes with higher lifespans in females but lower lifespans in males (Fig. 3c,d). This is due to a sex inversion in timing of sequential waves of mortality of carriers of the paternal haplotypes—early *D* mortality but late *H* mortality in females; early *H* mortality but late *D* mortality in males (Fig. 3f). *Vita2b* has a massive G×S effect with a LOD of 8.6 at T_140_ that is lost entirely by T_860_ (Fig.3a). The notable reversal of male genotype effects at *Vita2b* fits models of antagonistic pleiotropy (Fig. 3d) in that the polarity of effects on lifespan flip between the T_305_ and T_700_ survivorships—genotypes enhancing life expectancy in the first half of life that shorten life expectancies after 600 days.

*Vita2a, Vita2b,* and *Vita2c* are cautionary examples of how mapping both sexes together without including an interaction term can mislead. While the sex-combined analysis defines highly significant peaks at T_800_ for *Vita2b* and *Vita2c* (Fig. 3b) this is an artifact of not fitting the interaction term (Extended Data Fig. 1e,f). G×S mapping also unmasks two independent interactions at the extreme distal end of Chr 4 (154 Mb) at the supposedly male-only *Vita4b* locus: one at T_605_, the other at T_905_ (Fig. 3a).

The male-specific *VitaXa* locus encompasses the entire proximal 70 Mb of Chr X (Table 1, Fig. 3g). The *B* haplotype in the T_42_ survivorship confers a 30-day benefit compared to the *C* haplotype, and this effect erodes almost linearly with T-age. *VitaXa* is an example of an oligogenic locus that is likely to integrate effects of a number of recessive variants on this hemizygous chromosome in males. In heterozygous females the effects of *C* and *B* haplotypes do not differ in any survivorship (Extended Data 1ac, 2ac).

The profound sex difference in life expectancies in early survivorships (Fig. 1b) can be decomposed into sets of loci that specifically influence G×S interactions and early male mortality. The likely proximal cause is higher stress and aggression among co-housed males^57–60^. High male mortality is visible in all plots for *Vita1c*, *Vita2a*, and *Vita6b* (Fig. 2d, Extended Data 2, 3, 4). These and other loci have strong but transient effects earlier in life in males. The male disadvantage is eliminated in survivorships above T_725_ (Fig. 1b). From T_935_ to T_1100_ males typically are housed solo and gain a small 8-day advantage over females (T_1040_, *P* < .036). While studies of sex differences in longevity of mice have had mixed results^61^, there are unequivocal age-dependent sex differences in mortality rates in both wild^62^ and laboratory populations^18,19,63^. Sex differences in lifespan of humans have a different pattern: males consistently have 1.25 to 1.46-fold higher mortality rates than women in elderly cohorts without any convergence up to 100 years^64^.

### *Soma* loci direct trade-offs between body weight and lifespan

Mice were weighed at 42, 183, 365, 548, and 730 days. Weights at the first three ages correlate negatively with subsequent lifespans in both sexes (Fig. 5e) as expected from much previous work^1–8^. These rank order correlations differ greatly by sex—*ρ* values at T_185_ of –0.28 for males and –0.11 for females (*P* <0.001)^67^. This translates to a loss of about 14.3 days per gram in males and 3.7 d/g in females at the peak of reproductive performance (Fig. 5e,i,j). Sex differences remain significant from T_185_ to T_800_ (bold linkage score trace in Fig. 5e, FDR of 0.05 at a –log*P* of 1.64). Correlations in females are stable throughout adulthood with values close to –0.1 (Fig. 5e-h). In contrast, the negative correlation in males erodes with age and overlaps female values in the older survivorships (T_900_, Fig. 5f-h). Correlations become positive in both sexes after 730 days (Fig. 5h). We mapped body weights acquired at all five ages (±5 days) and detected 28 *Mass* loci that we subsequently used to compare with the *Soma* loci described next (Fig. 5a-c, Extended Data Fig. 7).

**Fig. 5.**
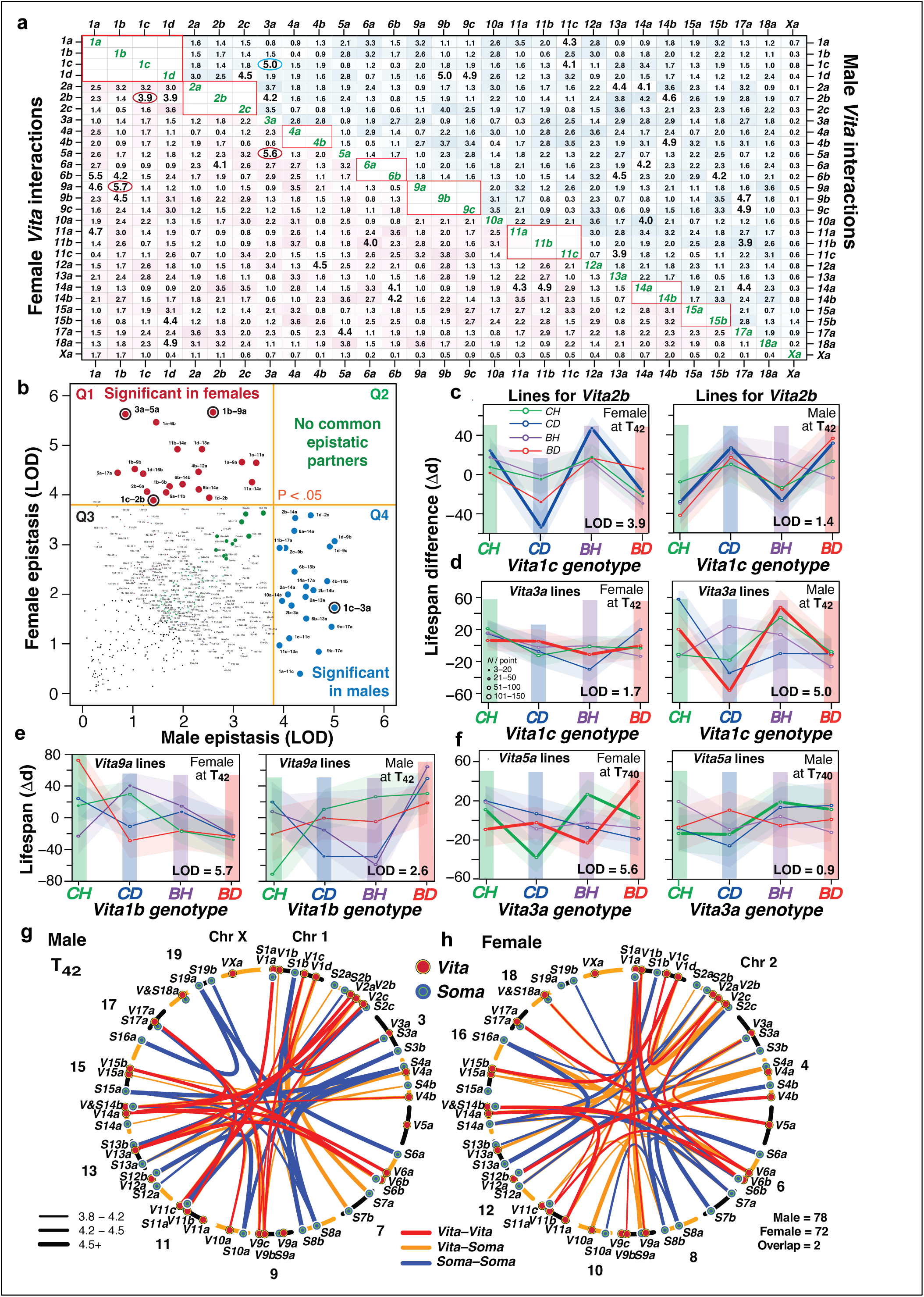
Epistasis among *Vita* and *Soma* loci. **a,** A full matrix of with LOD scores of the locus-by-locus interactions in the T42 survivorship of all animals in which the upper triangular matrix tabulates male scores, the lower tabulates female scores. LODs at or above 3.8 in larger bold font (*N* = 22 in males, 19 in females). There is minimal commonality between male and female epistasis. Similar matrices are given for all four ages for three sex combinations—combined, females only, and males only in Supplementary Data 10. **b,** Correlation plot of female versus male epistasis in the T_42_ survivorship. Those above threshold are marked in red for females (*N* = 19), in blue for males (*N* = 21) and labeled by *Vita* pair numbers. There are no points significant LODs shared by both sexes in quadrant Q2. However, the green points in Q3 highlight some low-level epistatic interactions shared by both sexes. The rho correlation of male and female LOD scores is 0.32. **c,** A pair of effect size plots of a strong female epistatic interaction between *Vita1c* and *Vita2b* at T_42_ that is also highlighted with a bullseye in **b**. The colored columns define the four genotypes at the first locus (*Vita1c*) whereas the four colored lines define genotypes of at the second locus (*Vita2b*). The *y*-axis is the lifespan difference of the 16 small points in each panel. Compare polarities of female and male effects of the *CD* genotype (thicker blue lines)—an example of modest antagonistic epistasis. **d,** A male interaction between *Vita1c* and *Vita3a* highlighted by the blue bullseye in (**b**). There is no interaction in females (left). **e,** Epistatic interactions at T42 and T740. The *Vita1b–Vita9a* interaction illustrates the classic masking effect with all lines converging in the pink *BD Vita1b* column. The male effects is not significant. **f,** The strong *Vita3a*-*Vita5a* interaction is weak in the T_42_ female survivorship but strong here at T_740_. Point sizes vary as a function of numbers of individuals per genotype pair. Almost all points are well above *N* = 50. **g,h,** Overview of all epistatic interactions in the T_42_ survivorship with LODs above 3.8 (thin lines), above 4.2 (medium lines), and above 4.5 (thick lines). Similar plots of the other three older survivorships are provided in Extended Data Fig. 9. Chromosomes are labeled with abbreviated *Vita* and *Soma* symbols. Color and type of lines define partnership types. There are 78 links between males and 72 in females of all types, but only two overlap in both sexes. Extended Data 9 provides the same plots for all survivorships.

To identify genetic causes of negative and positive weight-to-lifespan covariation we mapped genotype-specific differences in rank order correlations at all five ages across all applicable (later) survivorships—five actuarial scans in 15-day steps. We defined 30 *Soma* loci (Table 2, Fig. 5b,c, Extended Data 8) at which *ρ* differs significantly by genotype at one of the T-ages when weights were obtained (e.g., weight at 548 days tested at T_550_). Only seven *Soma* loci overlap the body weight *Mass* loci—a frequency expected by chance (*P* = 0.28, Fig. 5a vs. Fig. 5b,c, and Extended Data 7). Fifteen are detected only in males; four in females (Table 2). Nineteen *Soma* loci have significant *negative* correlations using body weights at 42 and 185 days. Eleven have significant *positive* correlations using body weights measured in the post-reproductive phase of life at 548 and 730 days^55,62^ (Table 2, Extended Data 8).

**Table 2.**
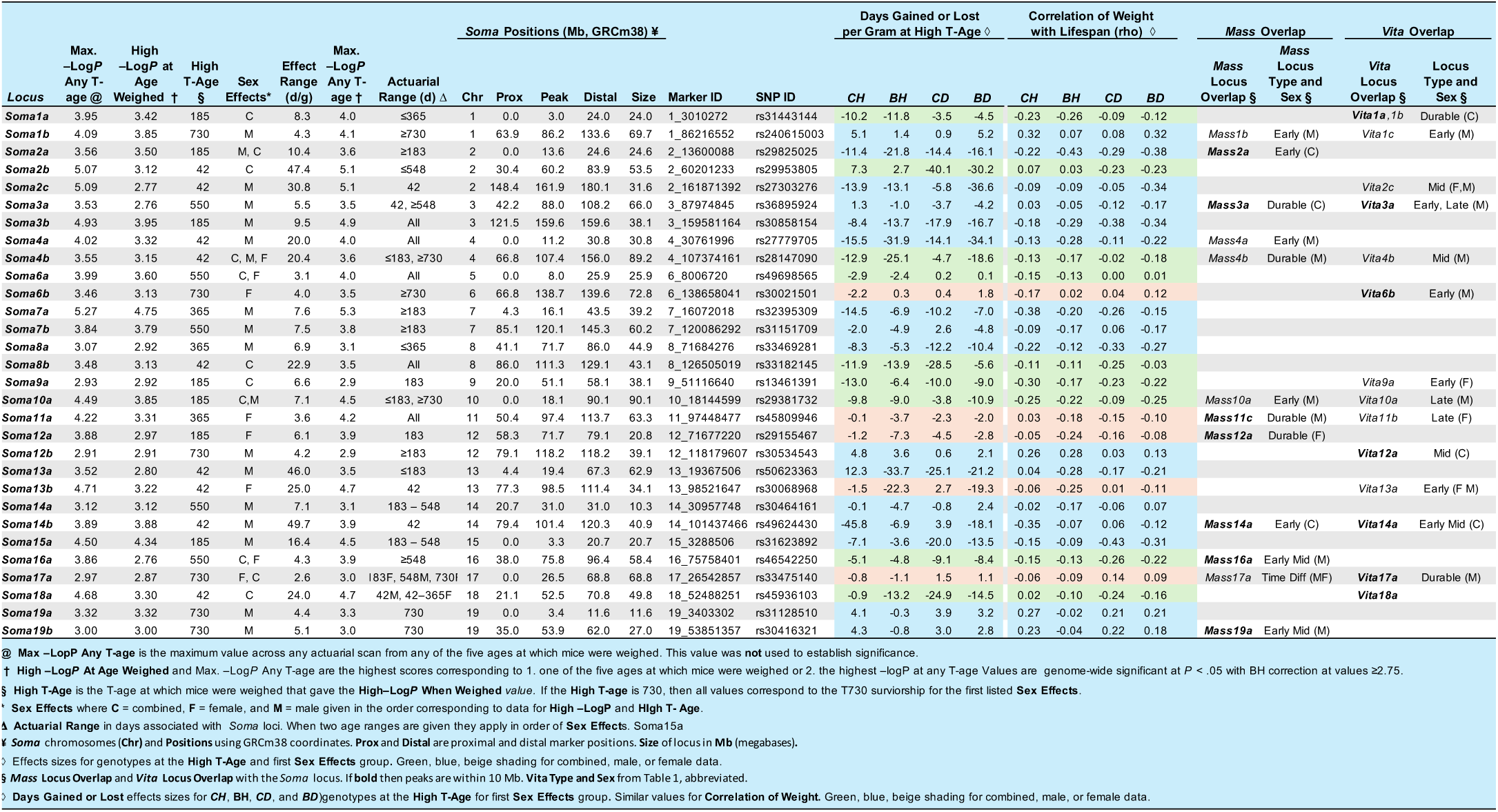
***Soma* loci modulate age-dependent correlations between body weight and lifespan**

The effects of *Soma* loci on life expectancies range from 2 days per gram to 29 d/g in T_42_, T_185_, T_365_, T_730_ survivorships (colored columns in Table 2). We highlight *Soma3b* that has strong effects in males (Fig. 5i) but none in females (Fig. 5j). Males with the *CD* genotype lose 17.9 d/g; those that inherit *CH* only lose 8.4 d/g.

In females, differences in negative correlations are insignificant (*CH:* –2.9 d/g versus *CD:* –3.4 d/g). *Soma11a* has the strongest impact on females (Table 2, Extended Data Fig. 5), but its effects—6 d/g between *CH* and *BH* genotypes—are comparatively modest compared to those of males.

*Soma* loci overlap *Vita* and *Mass* loci (Table 2) at a chance level. The summed genome-wide coverage of *Soma* loci is 1349 Mb or 48% of the mouse genome^68^, and coverage of the *Vita* loci is 1000 Mb (36%). Seven *Soma* loci have peaks within 10 Mb of *Vita* peaks, also insignificant (*P* = 0.19). But two *Soma* loci share the same peak marker with *Vita* loci, overlapping survivorship ages, and also have the same sex bias. *Soma1a* and *Vita1a* share top markers in both sexes, but the *Soma* effect is based on weight at 183 and 365 days (Fig. 5c,k,l) whereas the *Vita1a* effect endures to the T_900_ survivorship. *Soma1a* and *Vita1a* are both modulated by the *H* and *D* haplotypes (Fig. 1f, Extended Data 3a and 8a). *H* lengthens life by 12 days, while *D* shortens life by 22 days. This effect is shared by both sexes (sex-combined LOD of 6.5). Genetic effects of *Soma1a* oppose those of *Vita1a*, particularly in females. *H* decreases life expectancy by –4 d/g (average *ρ* = –0.12), whereas *D* increases expectancy by 1 d/g (*ρ* = +0.02). All effect plots, for example those in Fig. 5i-l, are provided in Extended Data 8 for all loci, both sexes, and T-ages.

### Epistasis interactions are segregated by sex and strongly age-dependent

We tested for epistatic interactions between both *Vita* and *Soma* loci at T_42_, T_365_, T_740_, and T_905_, stratified by sex. Fifty interactions are significant using the stringent Bonferroni correction for a total of 11,768 tests, but because we are most interested in patterns of epistasis, we analyzed all interactions with LOD ≥ 3.8—still significant at a Benjamini and Hochberg^69^ false discovery rate of <0.01. There are 41 *Vita*-*Vita* interactions among 359 *Vita*-*Vita* pairs tested in the base T_42_ survivorship with LOD scores ≥3.8 (Fig. 5a,b, Table 1)—19 in females and 22 in males. Across all four survivorships the number of unique *Vita-Vita* interactions is 43 in females and 59 in males (Supplementary Data 12.4). Similarly, there are 35 *Soma*-*Soma* interactions in females and 57 in males with LOD ≥3.8 (Fig. 5a,b, Table 1). Average LODs of interactions are 4.4 and 4.6 with standard deviations of 0.5 to 0.8. One of the stronger male interactions is between *Vita1c* and *Vita3a* (Fig. 5d, LOD 5.0) in which effects of *BD* and *CH* genotypes at *Vita3a* produce differences of 100 days in lifespan across the *Vita1c* genotype*s* (Fig. 5d). This interaction is lost by T_740_ (Fig. 5f). The strongest female *Vita-Vita* interactions are between *Vita1a–Vita6b*, *Vita3a*–*Vita5a,* and *Vita1b*–*Vita9a* (Fig. 5a,b). The *BD* genotype at *Vita1b* masks *any* effects of *Vita9a*—the classic Mendelian definition of epistasis (Fig. 5e)^70^. Finally, there are 197 *Vita-Soma* interactions—84 in females and 113 in males—a bias expected by significantly greater numbers of male *Vita* loci (20 vs. 8, Table 1) and *Soma* loci (18 vs. 8, Table 2). *Vita*-*Soma* loci also have mean LOD scores of about 4.5 ± 0.6 (s.d.).

Epistatic interactions of *Vita* and *Soma* loci form independent male and female networks (Fig. 5b,g,h, Extended Data 9). Even those few loci with minimal sex differences in their main effects do not share partnerships (Fig. 5a). For example, in females, *Vita1b* pairs up with *Vita6b*, *Vita9a*, and *Vita9b* with LODs of 4.2, 5.8, and 4.5, but in males LODs are negligible: 1.5, 2.6, and 1.1. No interactions are shared by *Vita* loci in the T_42_ survivorship (Fig. 5b), and only 2 of 61 possible interactions of all three types are shared at the most lenient threshold in any survivorship (Extended Data 9, Supplementary Data 12). The same pattern is true of *Soma*-*Soma* and *Vita*-*Soma* interactions. Polarities of interactions can also display sexual antagonism^71–74^, for example, the complementary effects of the *Vita2b CD* genotype (blue line) across *Vita1c* genotype columns (Fig. 5c). While *Vita1c* does not have any main effect in females (Table 1, Fig. 1e, Fig. 3a), the tests for epistasis expose a strong interaction in males of this locus with *Vita2b*^75^.

Epistatic partnerships are more stable in females than males as a function of age (Supplementary Data 12.3). For example, 21% and 17% of partnerships at T_42_ are also detected at T_740_ and T_905_ in females, but only 9% and 0% are detected in males. This is not caused by sex differences in strength of interactions. Essentially all of these findings are consistent with strong sex-dichotomized genetic architecture—a form of genomic diplomacy that fits male and female phenotypes with complementary life history goals^14,76–78^. These findings add to the list of genetic mechanisms that help accommodate two sexes into a single genome: 1. Sex chromosomes^79–81^, imprinting^82^, strong G×S interactions and apparent allele complementarity, and almost completely segregated epistatic interactions in the UM-HET3 population.

### From maps to mechanisms

Most *Vita* loci are too large for resolution of causal DNA variants or mechanistic dissection of mortality rates. However, 19,000 more UM-HET3 mice are available for genotyping, which already motivates mechanistic studies^25,46,83,84^. For each *Vita* locus we have assembled details on gene variants segregating in the UM-HET3 with predicted functional impact and roles in aging^85,86^. Here we highlight both limitations and prospects of analysis at this stage. The two largest loci almost certainly encompass many linked variants. *Vita17a* extends from the centromere to 74 Mb (Table 1), overlapping the major histocompatibility complex, known since the 1970s to be linked to lifespan^87,88^. *VitaXa* is detected only in males, and it covers the proximal 70 Mb of Chr X and is an intractable oligogenic locus. Males with the *B* haplotype live 30 to 15 days longer than those with *C* haplotype from T_300_ to T_1010_ (Fig. 3g)^89^.

In contrast, *Vita1a*, *Vita9b*, and *Vita15b* are tractable. *Vita1a* is compact, overlapping just 14 protein-coding genes of which *Atp6v1h*, *Rb1cca*, and *Mrpl15* are strong candidates involved in autophagy, nutrient sensing, and mitochondrial function^83^. *ATP6V1H* has ubiquitous activity as a modulator of vacuolar acidification^90^. It extends *Drosophila* lifespan^85,91^ and modulates insulin secretion and type-2 diabetes risk^92^. In the UM-HET3, the D haplotype of *Atp6v1h* also harbors an 11.5 Kb repeat in intron 7^93^. The IMPC homozygous knockout mouse is lethal early in life (www.mousephenotype.org/data/genes/MGI:1914864). Our finding that *Vita1a* has strong epistasis with *Vita6b*, *Vita9a*, and *Vita11a*, but only in females, is a strong constraint when evaluating gene candidacy using longitudinal multiomics data being generated for both UM-HET3 mice^94^ and BXD mice^95^. In preliminary mapping, this locus also modulates interactions of rapamycin with longevity using the same cases reported by the ITP in 2009^41^ with an interaction effect with rapamycin of LOD 4.5.

We end with a combined genetic and experimental analysis of *Vita9b*. This locus was mapped 23 years ago in an independent UM-HET3 cohort (Fig. 6a, Ref.^96^ their *D9Mit10* locus) and confirmed in the BXD family of mice (data from Refs^97,98^). Its replicability motivated us to perform a more intense dissection of genetic effects. *D* and *H* haplotypes reverse polarity in males between T_500_ and T_700_ and two strong inflections in late survivorships at T_845_ and T_1050_ (Fig. 6b, Extended Data Fig. 2o2). The *C* and *B* haplotypes have more uniform effects that peak at T_785_ (Fig. 6c). In females, effects are much more modest and delayed to T_800_ (Extended Data 2o2). These findings constrain and add useful priors for candidate gene selection.

**Fig. 6.**
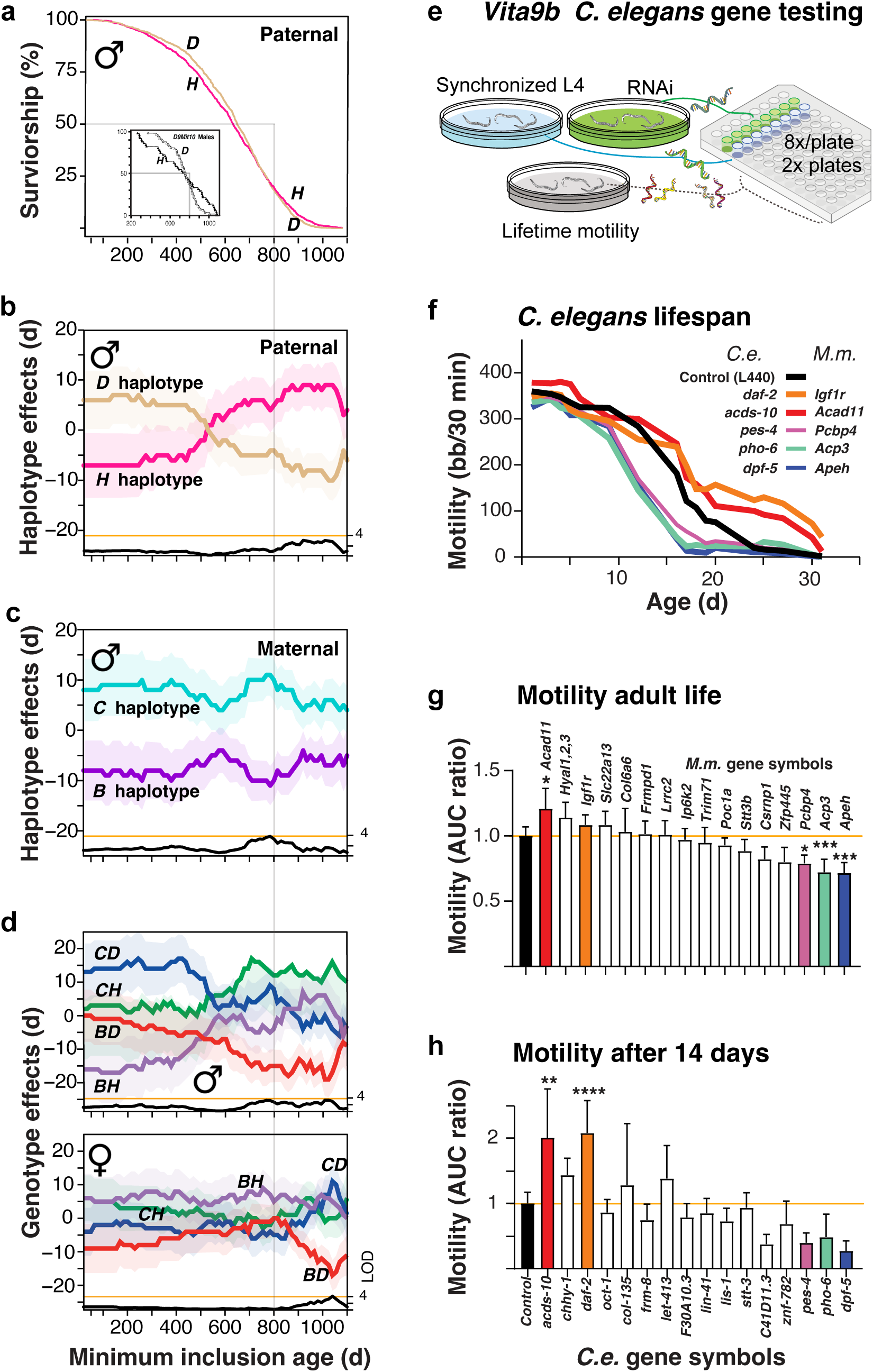
Genetics of *Vita9b* and candidate gene analysis. **a,** Kaplan-Meier plot for paternal haplotypes in male UM-HET3 mice at *Vita9b*. The *D* haplotype has a survival advantage compared to the *H* haplotype until 800 days (faint vertical line through panels **a-e**). The small KM insert box is redrawn from Jackson et al. (2002) who detected the same effect at this locus in a separate cohort (*D9Mit10*). **b,c**, Contrasting effect plots for paternal (*H, D*) and maternal (*C, B*) haplotypes along with LOD scores. Life expectancies of both *H* and *D* haplotypes invert between T250 and T900. Those of *C* and *B* haplotypes are durable. **d,** Corresponding genotype plots of males and females that are significant only in old survivorships. **e,** Design of the *C. elegans* motility assay in which RNAi were used to knock down expression of target transcripts with two levels of replication. **f.** Time course of age-related changes in motility of 6 of 15 lines—the control (black), and the *daf-2* positive control (orange). Effects of RNA knockdowns on motility of 15 orthologs of *Vita9b* candidates in *C. elegans*. **g,** Summary of motility over most of adult life (1 to 30 days). Asterisks are significant effects with correction for multiple tests. Of the 15 genes tested, 3 reduce motility significantly— *pes-4* (mouse *Pcbp4*), *pho-6* (*Acp3*), and *dpf-5* (*Apeh*). acds-10, an ortholog of the mouse *Acad11* gene, improves motility relative to control and in a pattern like that of *daf-2*. **h,** Motility after 14 days of age. Only effects of *daf-2* and *acds-10* are significant. All values are normalized to control integrated over the age range (ratios of areas under the curve; AUC).

*Vita9b* spans 127 protein-coding genes for which missense variants are predicted to have moderate or high impact. The region corresponds to human Chr 3p21.1 (46 to 53 Mb) and 3q22.2 (130 to 138 Mb)^99^. We tested 15 C. elegans orthologs using RNAi knockdown (Fig. 6e)—a validated lifespan predictor^100,101^. Four knock-downs significantly altered activity after 14 days (Fig. 6f). Suppression of acds-10, an ortholog of murine *Acad11*, boosted motility late in life—from 18 to 27 days—with an effect size approaching the *daf-2* positive (Fig. 6f-h)^102^. In mice, *Acad11* is trans-activated by p53, enhances fatty acid β-oxidation, and increases expression with age across tissues^103–105^. Three other knockdowns reduced activity starting after 10 days (Fig. 6f-h): orthologs of *ACP3* (*pho-1*)^106,107^, *PCBP4* (*pes-4*)^103–105^, and *APEH* (*dpf-5*)^108^. All candidates align with *Vita9b’s* late-onset action, but none have yet been linked to longevity^85^. Two caveats apply: candidate gene analysis is biased by the depth of the literature, and sparsely studied genes risk being deprecated—the so-called ignorome^109^; second, *Vita9b* is likely to contain multiple linked variants that contribute to late onset disease^110,111^.

## Discussion

Traditional genetics of aging has defaulted to a single metric—the duration of lifespan. In contrast, our method tracks mortality across nested survivorships from puberty to old age. This actuarial approach exposes the dynamics of mortality as a function of genotype, age range, and sex. At least 59 loci *Vita* and *Soma* loci shape mortality risks and lifespans of UM-HET3 mice. Some act early, others late, and most differ greatly in their timing and their impact on males and females. These loci participate in hundreds of epistatic interactions that are segregated almost completely by sex. A subset of loci have persistent genetic effects on mortality rates from T_215_ to very old survivorships and may be rare modulators of rates of aging. Males have many early acting *Vita* and *Soma* loci, and about 25% more epistatic interactions than females across all survivorships. Doubling our previous sample size revealed complex patterns of interactions and confirmed six of seven loci we initially detected^46^. Knowing when genetic effects appear and disappear provides key context for molecular and cellular networks that alter age-dependent mortality risks across lifespans^10,112–114^.

The UM-HET3 mice are a single sibship derived from a cross of isogenic F1 hybrids, and like humans they segregate for millions of sequence variants. Unlike humans, UM-HET3 mice have a uniform and very high minor allele frequency of 0.25—a benefit in terms of statistical power that contributed to our success in detecting loci even in the oldest 20% of survivors (*N* = 1568). This is equivalent to studying the genetics of longevity in those humans who have reached their 90s^115,116^. We are able to control diet and environment tightly and thereby minimize many types of confounds—in particular, infectious diseases and climate. A disadvantage of this murine family is that loci cannot yet be mapped with high precision, although two *Vita* loci were bracketed into 4–8 Mb intervals.

### The age-dependence of genetic effects on lifespan and mortality

We listed four questions in the introduction. The first relates to the age-dependent effects of loci that modulate survival. Some such as *Vita1c* and *Vita18a* act on mortality rates only early in life with effects lost after 500 days. These loci are much more common in males than females and are likely to be linked to male-male competition, hormone status, and stress resilience^67,117^. Other loci act almost exclusively from 500 to 700 days (*Vita4a*, *Vita11a*, *Vita12a*). A final set of loci affect only the oldest-old individuals (*Vita5a*, *Vita9b*, *Vita9c*). Our unsurprisingly answer to this first question is that loci that influence mortality rates have strong age-dependent effects, as do their interactions. The actuarial approach, combined with a two-fold increase in sample size, greatly boosted the number of loci we have been able to detect and places them securely into age-dependent classes that fit well with evolutionary and life history theories of aging.

### On genetics of lifespan and evolutionary theories of aging

Our second question was whether different types of age-dependent effect on mortality support evolutionary explanations of aging^36,38,39,50^. The first theory is that aging is caused by late acting variants that escape selection in natural populations due to high extrinsic causes of mortality—Haldane and Medawar’s *mutation accumulation theory*^36,37^. This idea predicts a range of post-reproductive genetic effects—both good and bad with respect to longevity, that do not have detectable effects on mortality early in life. In mice their impact on mortality should emerge only after 500 days; either gradually like *Vita5a* in males or abruptly like *Vita1a* in females. The second theory is that aging is a consequence of a fitness optimization and a careful compromise between reproductive investment and the necessity of self-maintenance and repair^39,50,118^. This is Kirkwood’s *disposable soma* theory^39^. The *Soma* loci collectively support this theory— they fit the model of loci mediating a bioenergetic compromise between size and reproduction and duration of life that are common in *r* selected prey species adapted to volatile environments^119,120^. The third theory is that aging represents a Faustian bargain between DNA variants that boost fitness early in life, but call for payment late in life—Williams’ *antagonistic pleiotropy*^2,38,121^. Antagonistic pleiotropy is vivid in loci such as *Vita2a*, *Vita2b*, *Vita2c*, *Vita9b* (Fig. 6b) with effect polarities that invert with age. A simple prediction is that alleles that improve fitness before 400–600 days will be linked to higher reproductive success but increased late-life mortality—a pattern that is much stronger in males than females. The summary answer is that the dynamics of *Vita* and *Soma* loci provide a firm empirical bridge between theories and organismal processes of aging^2,113,122–124^.

### On sex and genetics of aging

Our third question addressed sex differences in lifespan. In our previous work^125^ we detected one female-specific locus on Chr 3 using the full survivorship and found no significant male-specific loci or gene-by-sex interaction effects, a limitation we attributed to low statistical power^46^. By doubling sample size and using actuarial mapping we now define 13 loci with their strongest effects in males, 4 with their strongest effects in females, and 12 loci with their strongest effects when data are pooled across both sexes (Table 1). This male bias is just as pronounced among the *Soma* loci—16 in males, 5 in females, and 9 combined. Sex differences in lifespan genetics are further accentuated by a deeper analysis of epistatic interactions of these loci, mirroring findings in *Drosophila melanogaster*^5,126^. Sex differences can be so divergent that we use the labels *sexual antagonism* or *reciprocity*. Of course, this imprint of sexual competition in much stronger during the reproductive crescendo of life, contributing to trade-offs between body mass and life expectancy that are highlighted by the male *Soma* loci. After reproduction, male and female life expectancies converge—natural selection has already had its say.

### The integration of *Soma*, *Mass*, and *Vita* loci

Our final question focused on trade-offs between body size and life expectancies. We have converted the inverse correlation between weight and lifespan into discrete *Soma* loci with strong sex- and age-dependent effects. At one level the *Soma* and *Vita* loci appear to modulate expectancies autonomously, but at the level of epistatic interactions they are strongly intertwined. And while 12 *Soma* loci overlap the *Mass* loci that control body weight, their time courses and sex effects rarely align (Fig. 5, Extended Data Fig. 5). The *Soma* loci directly exemplify antagonistic pleiotropy, defined here as modulators of the trade-off between body weight and lifespan. These are the first such loci mapped in vertebrates with clarity. They recapitulate the equally explicit trade-offs in *Drosophila*^123,124,126^. Why is there such a marked sex difference in the genetic architecture of *Soma* loci? We propose the same explanation as for *Vita* loci; that differential selection pressures favor fast growth and physical robustness or slow growth with better resilience and survival under stress and resource limitations^62,127^. The independence of most *Soma* loci from *Vita* and body *Mass* loci demonstrates that distinct biological processes orchestrate the mass-mortality equation in males and females.

#### Candidate gene analysis and the benefits of delayed gratification

Variation in the dynamics of *Vita* and *Soma* loci likely stems from allelic differences affecting growth rates, fitness, fecundity, and functional and genomic robustness across ages, and inevitably, the effects of these differences on disease susceptibility and onset. Wild mice die mainly of predation, disease, and thermal stress^127,128^ while laboratory mice die mainly of cancers and, less commonly, from renal, cardiovascular, and inflammatory diseases^97,129,130^. Given the many intrinsic and extrinsic factors that are in-play and the large size of most loci we expect loci to blend the effects of linked polymorphisms^131–133^. The main challenge of candidate gene analysis of complex traits is when to say “not yet”. In this work, we have tried to outline our approach to this problem and to balance the promise against the risks of premature analysis. While the oligogenic *Vita17a* and *VitaXa* loci are out of scope, *Vita1a* and *Vita9b* merit attention. Several candidates—*ATP6V1H*, *ACAD11*, *ACP3*, *PCBP4*, and *APEH*—may already justify *in vitro* and *in vivo* tests and dissection of the phenotypes and pathologies of individuals with defined genotype classes over specific age ranges.

#### The dynamics of heritability of variation in life expectancy in mice and humans

Genetic factors account for roughly 25% of variance in life expectancy of young adult humans^7,134^, but heritability appears to increase as a function of age^35^. This trend has motivated searches for DNA variants in the oldest old humans^21,24,135^. In UM-HET3 mice we also find evidence of increasing heritability with age in males but less so in females (Fig. 2g,h). Surprisingly, in the oldest-old mice there is a decline in environmental variance, perhaps associated with the loss of lifelong companions or higher order dysregulated genetic effects. Does it make sense to focus on gene variants acting principally in the oldest-old? We do not think so. They are happenstances of the shadow of selection. Far more important are those loci that modulate mortality from early stages of life through to senescence—the rate of aging modulators (Table 1)^47,136^. We highlighted 14 *Vita* loci of this type that may harbor variants that are both selectable and controllable.

#### Prospects

*Vita* and *Soma* loci can obviously be used to predict lifespan of UM-HET3 mice, but we have not done so because our maps and models need further refinements. Higher density genotyping of the additional 19,000 UM-HET3 mice—including those treated with supplements such as rapamycin that prolong lifespan— will provide us a much more secure foundation for prediction. Models will also need to be fine-tuned for environment, diet, sex, maternal age, litter size, parity, and higher order terms such as three-loci interaction effects and social genetic effects among co-housed mice^56,137^. ITP teams have spent 20 years rigorously testing 74 different compounds for their lifespan-extending potential. Eleven of these drugs have had significant positive effects—an impressive 15% hit rate^40–42,138–146^. But less noted but of prime importance, there has been an unusually strong sex bias in favor of positive effects in males (updated through to 2021 cohort). Rapamycin is one welcome exception with strong beneficial prolongevity effects in both sexes—but this drug like all others will be strongly modulated by many gene-by-drug interactions. One person’s poison is another’s cure and sex age and many genetic differences will obviously matter. We have shown here that all ITP drugs can now be placed safely within a known constellation of 59 loci and potentially hundreds of epistatic interactions that modulate mortality, potentially guiding interventions to extend human health and longevity^147–151^.

## Materials & Methods

### The UM-HET3 sibship

UM-HET3 mice are progeny of a cross between two types of F1 hybrids—female F1s from matings of BALB/cByJ (C) dams to C57BL/6J (B) sires, and male F1s from matings of C3H/HeJ (H) dams and DBA/2J (D) sires. These four inbred progenitors are abbreviated CBy, B6, C3H, and D2 when referring to mice and strains, and abbreviated *C*, *B*, *H*, and *D* in italic when referring to genotypes and haplotypes. They were selected to maximize phenotypic diversity. Young virgin F1 males and females were distributed from the JAX production facility at The Jackson Laboratory (TJL) to ITP research colonies at TJL in Bar Harbor Maine (JL), the University of Michigan in Ann Arbor Michigan (UM), and the University of Texas Health Science Center in San Antonio (UT). Breeding cages were set up each spring. First litters were not used. All subsequent litters were used at UT, litters with 6 or more pups were used at JL, and those with 7 or more were used at UM. Weanlings were entered into the study over the next seven months. Mice lived together from weaning at 19 to 21 days and without replacements until their deaths^152^. As a result, most mice live solo after 1000 days, a factor to consider with respect to mortality risks but not yet integrated into the analysis here.

All mice in the first four cohort years were born between April 2004 and January 2008. For these years we have numerically well-balanced DNA samples from all sites. Almost no animals were generated in 2008 due to a funding gap. All mice in the final cohort years used in this study were born between July 2009 and March 2013. The 2009 cohort includes mice from all sites, but 2010 and 2011 include mice only from UM and UT. Cohort years 2012 and 2013 include mice only from UM. The last UM-HET3 mouse in this studied died on Dec. 21, 2015. In all years, only a very small percentage of mice (<10%) were born from January through April. Mice were weighed at 42 ± 2 days, and at 183 days (6 months), 365 days (12 m), 584 days (18 m), and at 730 days (24 m) with a timing error of approximately 7 days. In all years, only a very small percentage of mice (<10%) were born from January through April. Mice were weighed at 42 ± 2 days, and at 183 days (6 months), 365 days (12 m), 584 days (18 m), and at 730 days (24 m) with a timing error of approximately 7 days.

The UM-HET3 sibship segregates for ∼10.6 million sequence variants (Supplementary Table 1 zip-file). All UM-HET3 mice inherit *C* and *B* haplotypes from their F1 mothers and *H* and *D* haplotypes from their fathers. As a result, sibs segregated for four genotypes (four two-way combinations of maternal and paternal haplotypes) on all autosomes—*CH*, *CD*, *BH*, and *BD.* Females inherit one entire non-recombinant *H*-type Chr X from their paternal grandmothers and a potentially recombined Chr X from their mothers (recombinations between *C* and *B* haplotypes only, see Fig. 3g). As a result, females have either *CH* or *BH* genotypes. Hemizygous males inherit a potentially recombined *C* or *B* Chr X from their mothers. All mitochondria are derived from maternal grandmothers (*C*) and Chr Y is derived from paternal grandfathers (*D*).

We genotyped 6,872 UM-HET3 mice for which we had full lifespan estimates (*N* = 3252 females, *N* = 3620 males). Of these genetic siblings 6,438 passed all genotype QC steps at 893 high-quality markers (*N* = 3401 females, *N* = 3037 males; see *Genotyping and Marker QC*). To ensure balanced numbers by sex later in life, every ITP cohort initially consists of 51 male and 44 female weanlings per year, per site, and treatment category, but with two-fold more common untreated controls at each site. This was done to enable overly aggressive males and any wounded animals to be removed while balancing male and female numbers later in life— almost always before 550 days. The earliest minimum inclusion age in this study is T_42_, the pubescent age at which tails were docked for tissue acquisition, an age equivalent to about 12 years in humans^153^. For numerical convenience we set the first truncation age at T_35_. To compensate for two-fold higher male mortality in the first two years of life (33% vs 16%) we included 12% more males than females in the T_42_ survivorship. The earliest death was at 46 days. This left us with a small surplus of 227 males at T_365_, but by T_560_ numerical balance was restored and there were 2930 females and 2929 males. This age corresponds to roughly 56 years of age in humans (Ref.^154^ their fig. 20.3). All individual-level data, metadata, and survivorships, and mortalities per 15-day interval are provided in Supplementary Table 2.

We genotyped two major categories of mice—those not treated with any dietary intervention and mice treated with a dietary supplement that did not modify lifespan significantly on a per-drug basis using standard statistical criteria^138,142,155^. These latter mice have been referred to as *No Drug Effect* (NDE) cases. However, when data are combined across the entire class of these individually “ineffective” agents from 2004 to 2013, there is in fact a highly significant multi-supplement that is positive on lifespan (Results). A better term going forward is *No Significant Effect* (NSE).

### Husbandry

Mice were weaned into same-sex cages—three males or four females per cage—at 20 ± 1 days^40^. From 2004 until 2013, all sites used NIH-31 standard diets. For breeding cages, UM used Purina 5008, UT used Teklad 7912, and JAX used Purina 5K52. For mice up to ∼122 days, UM used Purina 5001, UT used Teklad 7912, and JAX used Purina 5LG6. After 2004, the control diet LabDiet 5LG6 was used by all sites. Mice were monitored daily for signs of ill health and aggression. Mice were euthanized if moribund. Tails were obtained at 42 ± 2 days for DNA extraction. Body weight at this age was acquired for 2459 mice, and at half-year intervals for 4688 mice at 183 days (6 months), down to 2208 mice at 730 days (24 m). Our analysis includes four experimental variables—sex, site, dietary drug treatment, and cohort year (Supplementary Table 3). Treatments are listed on *Mouse Phenome Database ITP Portal* under the acronyms 4OHPBN, CAPEhi, CAPElo, Cur, Enal, FOhi, FOlo, GTE, HBX, I767d, MB, MCTO, MET, NFP, OAA, Res07, Reshi3, Reslo3, Simhi, Simlo, and UA.

### DNA extraction from tails, sequencing and sequence alignment

Tail tissue was processed using the MagMAX magnetic-bead extraction system and genomic DNA was quantified using a Qubit 4 fluorometer and a high-sensitivity DNA assay kit (Thermo Fisher Scientific). Locus-specific PCR primers were designed using *BatchPrimer3*^156^ to generate 180 bp amplicons. They were mixed using an optimized Hi-Plex approach at Floodlight Genomics (Knoxville, TN). Sample-specific 12 bp barcodes were added and amplicons were pooled^157^. Primer sequences for SNP markers with rs-type identifiers are provided in Supplementary Table 4). The amplicons were sequenced to an average of 1000X per targeted locus on an Illumina NovaSEQ (2x150 bp). Data were demultiplexed and FASTQ files used for genotyping. FASTQ files were aligned to *Mus musculus* GRCm38.p6/mm10 reference using *Bowtie 2* (version 2.3.4.1)^158^. BAM files were sorted and indexed using *samtools* (version 1.6)^159^, and read group information was added using *picard* tools (version 2.14.1).

### Variant calling

We called short sequence variants using *bcftools*^159^ in three steps: 1. we created text pileup output for all BAM with *mpileup* at a mapping quality of 30; 2. we called using default settings; 3. we removed variants with a read depth <100 across all samples and a QUAL score <100. Sequencing depth per case was used as an additional criterion for filtering. Sequence for C57BL/6J, C3H/He, and DBA/2J was downloaded from the Wellcome Sanger Mouse Genome Project^160^ (www.sanger.ac.uk/data/mouse-genomes-project) and that for BALB/cByJ was generated by Beierle and colleagues^161^. From these files we extracted variants that differ among founders (Supplementary Table 1, a large *zip* file). Variants were called in three steps as above, the differences being that the minimum depth was reduced to 10x and QUAL score to ≥30. Variants that were called reliably in all founders were advanced for genotype phasing and mapping. VCF files with variant data on UM-HET3 and their founders are provided in Supplementary Table 5.

### Genotyping by Sequenom MassARRAY

Tails were plated in deep-well plates and sent in three different batches to Neogen Corporation (Lincoln NE). Individuals were genotyped by SNP genotyping by Sequenom MassARRAY MALDI-TOF mass spectrometry^162^ at a maximum of 270 markers. Fifty markers were removed after QC, based on deviations from expected allele frequency (0.35 < frequency < 0.65). We chose pairs of linked markers to define maternal and paternal chromosome genotypes.

### Phasing SNP genotypes

SNP genotypes were phased using R v4.0.2 (*R Development Core Team* 2005; script at *github.com/DannyAr-ends/UM-HET3*). We generated fully phased haplotypes for all sets of maternal and paternal autosomes and Chr X. Markers fall into those that unambiguously define maternal haplotypes (*N* = 486)—*C* vs *B* alleles (denoted by lines in pink Fig. 1c); those that define paternal haplotypes (*N* = 396)—*H* vs *D* alleles (blue lines in Fig. 1c). We generated diploid genotype probabilities at all positions from phased chromosomes using the *calc.genoprob()* function in R/qtl. Markers were reviewed and those that had a call rate <30% were excluded. When we plot genotype and/or haplotype effects (Extended Data Figs 1 and 2), we call genotypes and haplotypes when imputation certainty is greater than 80%. We used R/qtl to estimate number of recombinations per mouse. The mean recombination number was 25.1 ± 8.6 SD. Just over 370 animals had recombination numbers more than two standard deviations above this mean. We excluded 59 mice that had more than 80 recombinations—likely to be caused by sample cross-contamination. The final genotype files are available at *GeneNet*work.org^163–165^ by setting *Species* = Mouse, *Group* = Aging Mouse Lifespan Studies (NIA UM-HET3), and *Type* = DNA Markers and SNPs and pressing the *Info* button. Supplementary File 6a lists all genotypes, and Fig. 6b is the R/qtl cross object used for most analysis.

### Genotype quality control

Markers and individuals with greater than 90% missing data were removed from further analysis (*N* = 333). We converted data from genotypes into parental haplotypes (see section: *Phasing SNP Genotype*s). Markers were checked for Mendelian inheritance errors and removed when the missing the data rate after conversion exceeded 90%. We integrated MonsterPlex and Sequenom marker sets based on genomic positions. We inspected the marker map for linkage between neighboring markers, and removed a single marker (Chr 4_58784823) which did not show linkage to its neighbors. We also removed 20 individuals that died of unnatural causes and 22 that we inadvertently genotyped that had received an effective drug supplement. Finally, we constructed an R/qtl cross-object for imputation using the remaining mice, imputed their haplotypes based on the full population (*N* = 6,438) (Supplementary Table 7).

### Inversions in the UM-HET3

There is a large inversion on Chr 6 of the C3H/HeJ progenitor strain^166^ between 51 and 94 Mb and may affect map precision (Table 1, 2). The recombination fraction in this interval is 0.00025 versus an expectation of 0.21. At least 42 Mb of the paternal chromosome is locked in linkage. We detected a total of 13 potential inversions ranging in size of 3.5 to 54 Mb. Six suppress recombination on the maternal chromosome and 7 suppress recombination on the paternal chromosome (Supplementary Table 8). There is overlap of maternal and paternal inversions on Chrs 4, 7, and 11.

### Lifespan estimates depend on age of entry into a cohort

Based on work by König and colleagues^62,127^ we know that in wild commensal populations of mice, 35–40% of pups do not survive beyond weaning. At birth, mean lifespan can be as low as 196 days. By 13 days, the mean expectancy has improved to 250 days for males and 500 days for females. We make these points to emphasize that lifespan estimates are made with reference to an operational starting point—a minimum ascertainment base age for a population. Researchers often incorrectly imply that mean lifespan is estimated from the date of birth, but this neglects earlier and often unknown numbers of deaths. In our case we can only measure lifespan for UM-HET3 progeny that lived to be pubescent juveniles. When we say that the mean lifespan of the 6438 sibs in this study is 844 ± 210 SD days (median of 870 d), we mean that this is the mean age at death conditioned on a mouse having lived to at least 42 ± 2 days—the age at which tail tips were taken for DNA extraction.

In a similar way we can determine the mean lifespan or life expectancy of the survivorship subsets of UM-HET3 mice that lived to be at least 185, 365, 545, 725, or 1100 days (roughly 0.5, 1, 1.5, 2, and 3 years, respectively). The mean conditional lifespans of those five survivorships or age cohorts are 847 ± 206 SD days, 861 ± 188, 890 ± 160, 941 ± 126, and 1162 ± 54 SD days. The oldest mouse was a female that died at 1456 days. We refer to these different ages at which lifespan can be computed as either survivorships or cohorts.

The actuarial procedure consists of sequentially studying progressive older and smaller survivorships from the radix population of 6438 mice in Fig. 1a. The minimum inclusion age is 42 days. Between T_42_ and T_50_ days a single male died. We truncated the survivorships upward from T_50_ to T_1100_, in 15-day steps. Although all individuals have accurately known lifespans, we sometimes use the term *life expectancy* to emphasize that our approach exploits lifespan data in an actuarial context conditioned on having reached a specified age. We speak of the life expectancy of mice that have reached 1100 days in the same way that we would speak of the life expectancy, not the lifespan, of 70-year-old humans.

The actuarial mapping strategy is affected both by lower sample size and by right skew at higher truncations. To ensure that our results are not affected by this, we used three approaches. First, we use a non-parametric quantile regression in combination with standard parametric mapping to make sure associations are detected using both methods^167^. Second, we limited ourselves to a maximum truncation age of T_1100_. Third, we used a time-dependent hazard function to validate *Vita* loci.

We tested a complementary approach for mapping under the assumption that loci contribute to variation in the time-dependent hazard function as in our previous^125^ but with refinements. We carried out a test at each marker of the null hypothesis that the (time-dependent) hazard function of death does not depend on genotypes. To do this, we used a model where the hazard function is allowed to depend smoothly on age with haplotype and baseline covariates, using natural splines with three degrees of freedom. For computational tractability we created risk sets with 50-day window^168^.

### Implementation of actuarial mapping

We mapped in 15-day survivorship steps from T_42_ to T_1100_. Mapping was performed using both conventional four-way mapping of genotypes at 893 markers (Fig. 1c) or by restricting analysis to each of the parental haplotypes—*C* versus *B* on the maternal chromosomes (*N* = 497 markers), and *H* versus *D* on the paternal chromosomes (*N* = 396 markers). Note that Chr X only segregates for maternal *C* and *B* haplotypes in both sexes (Fig. 3g). In general, mapping all four genotypes provides better power. This method defined all loci in Table 1 and full LOD scores across all survivorships are illustrated in Extended Data Figs. 1, 2. The null model (model(H_0_)) and alternative model (model(H_alt_)) were fitted at each marker using the following linear models:

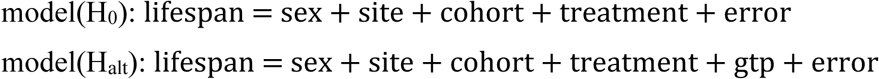

*Sex* has two levels (M, F), *site* has three levels (JL, UT, UM), *cohort* year has nine levels (no animals were generated in 2008), *treatment* has two levels—untreated controls and cases that received a drug that had no significant effect on lifespan. The gtp (***g***eno***t***ype ***p***robability) term is a *N* row by four-column matrix, where *N* is the number of individuals used in the mapping, and columns are genotype probabilities for markers computed by R/qtl (v 1.48-1)^169,170^. Sex was dropped as a co-factor from the regression model when mapping was stratified by sex. LOD scores were computed by comparing the fit of the null model with the fit of the alternative:

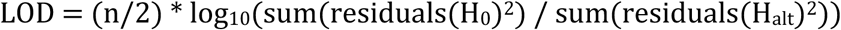

Mapping on Chr X requires special care due to the four-way cross structure and greater difficulty defining highly reliable markers for haplotype, particularly close to both telomeres.

### Effect size plots resolve survivorship differences by genotype and sex

We use the four-way mapping algorithm implemented in R/qtl^170,171^ while controlling for sex, site, cohort year, and drug treatment. We compute maps for survivorships starting from T_42_ up to T_1100_ (Fig. 1a,d,e,f). We introduce a new type of actuarial plots in Fig. 1b,d,e. The *x*-axis defines the minimum inclusion age (T-age) of each survivorship. The expected increase in error terms in older survivorship with lower sample sizes are mitigated by the reduced range over which animals die within progressively older survivorships—from 1410 days at T_42_ to 356 days at T_1100_. Errors of lifespans of survivorships are relatively stable—from 844 days ± 210 SD in the T_42_ base population (SD/ range of 15%) to 1162 ± 54.4 SD in the last T_1100_ survivorship (SD/range of 15%). When interpreting effect size plots note that increases and decreases in mortality rates among subgroups are relative to the mean lifespan of that T-age.

The T-age of a survivorship always refers to an entire age *range*. Actuarial truncation in a forward direction means that the oldest-old mice are embedded in every survivorship. Reverse truncation flips the polarity by truncating from oldest to youngest and defines *exitships*. The _1099_T exitship consists of mice that died before 1100 days and complements the T_1100_ survivorship. The combination of forward, reverse, and even two-sided truncations is useful to dissect causes of mortality in age-restricted subpopulations. In this paper we restrict attention to forward truncations.

### Significance testing for actuarial time-series analysis

We evaluated the significance of mapping results across two axes: 1. the spatial location of markers and their independence across the genome; and 2. the temporal axis and the number of effectively independent actuarial tests. We mapped using 891 markers but given the significant linkage between markers we effectively tested about 442 independent genome locations. This number of independent tests was estimated using the *simpleM* method with a block length ranging from 10 to 500 markers and a 10-marker step size^172^. The threshold at a genome-wide type 1 Bonferroni error rate of 10% is 3.65 LOD (–log_10_(0.1 / 442), at 5% it is 3.95, and at 1% is 4.65. These are conservative thresholds. For maternal and paternal maps, the density of recombination is reduced by half, and corresponding thresholds are 3.44, 3.65, and 4.34.

Positional confidence interval using the typical 1.5 LOD-drop in linkage^51^. We also estimated temporal range of action of loci, defined as the T-ages over which a locus has strong effects. The start and stop T-ages are those that immediately precede or succeed survivorships that are genome-wide significant. For example, if linkage is above LOD of 3.95 from T_365_ to T_725_, then the T-age CI of the locus from T_350_ to T_740_. This criterion is not conservative when only a few survivorships have significance. In this latter case, we define the T-age range that brackets the peak T-age by a >1.5 LOD drop.

We evaluated the impact on significance testing of mapping multiple nested survivorships using a Cauchy combination test^173^ at each marker across all survivorships (Table 1). Cauchy *P* values were adjusted for multiple testing using a Benjamini and Hochberg correction to account for multiple testing across markers, and the Cauchy –log*P* in Table 1 of ≥1.50 is significant at a *P* of less than 5%.

### Locus assignment and candidate gene region determination

Locus names are prefixed with *Vita*, *Soma*, and *Mass* identifiers followed by chromosome and a letter suffix (*Vita1a*, *Soma2b*, *Mass11c*). When the 95% confidence intervals of two adjacent QTLs overlap, and the genotype and haplotype effects appear to be the same, loci are generally assigned the same *Vita* identifier. For such “merged loci”, the 95% CI is reduced to the overlapping interval. Determining whether a locus is detected in different subsets of the cohort is somewhat subjective, examples being *Vita3a* and *Vita14b* that could have been split into two loci by sex and T-age. We provide data for linkage scores in all three categories (sexes combined, females, and males) for *Vita* loci (Supplementary Table 9), *Soma* loci (Supplementary Table 10ab), and *Mass* loci in (Supplementary Table 11).

### Testing for sex interactions at *Vita* loci

There are significant differences between male and female stratified survivorship maps. To investigate this formally and to detect significant sex-marker interactions modulating age-specific lifespan we tested sex-interactions of *Vita* loci at the T-age with the peak LOD (Fig 3a). We computed the LOD score difference of two models against each other:

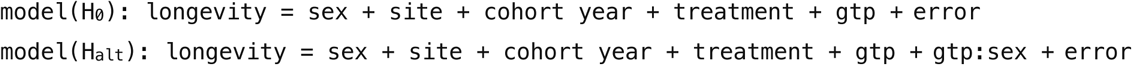

Since we are only interested in significant sex-interactions at previously detected *Vita* loci but excluding *VitaXa*. We used a Bonferroni correction based on 28 markers. A LOD scores above 2.75 is significant (– log_10_(0.05/ 28)).

### Testing pair-wise epistatic interactions between *Vita* loci, *Soma* loci, and *Vita*-with-*Soma* loci

A two-dimensional analysis of marker-marker interactions affecting life expectancies was performed by testing the following models against each other in four survivorships—T_42_, T_365_, T_740_ and T_905_.

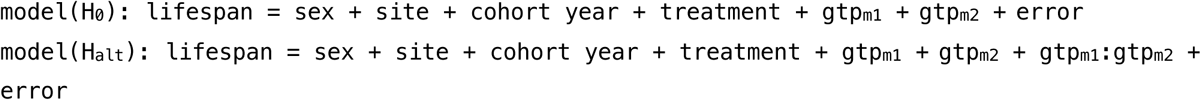

We computed the difference in fit between H_0_ and H_alt_ as explained in the section “Implementation of actuarial mapping”. To limit numbers of tests, we initially tested interactions between the 29 *Vita* loci (Table 1), excluding all pairs of loci on the same chromosome (Fig. 5a). This resulted in the matrix of 359 non-syntenic tests of pair of loci per sex. We used Bonferroni corrections for most work. With 359 *Vita*-*Vita* tests a LOD of about 3.8 is appropriate, but for more comprehensive tests of all loci in all four survivorships (11,768 tests) the corresponding *P* <0.05 threshold is 5.37, a value that limits the yield of interactions to only 50 (Supplementary Table 12.2). In contrast, the Benjamini and Hochberg test suggests that 966 interactions have FDR of less than ≤0.005 (Supplementary Table 12.2). We opted for a compromise and analyzed a total 289 unique epistatic interactions that are visualized in part in Fig. 5g,h for the T_42_ survivorship and in full in Extended Data 9 for all four survivorships. All of these interactions have LODs of ≥3.8. Of the total of 11,768 tests fewer than 4% have LODs ≥ 3.8 (Supplementary Data 12.)

### Visualizing the genetic modulation of mortality rates

We plotted age-dependent mortality and the hazard ratios (HR) of haplotypes to give more insight into the specific ages over which genetic differences operate with more or less force. We computed mortality rates of maternal and paternal haplotype pairs (*C* versus *B*, *H* versus *D*) for males and females separately. We used LOESS smoothers with span parameters of 0.2 (Extended Data 4), and adaptive LOESS with a span of 0.3^174–176^ with shorter spans over ranges with high numbers of deaths (800–950 days) and longer spans over ranges with few deaths (Fig. 2f). Log2 HR ratios greater than 0.6 represent relatively strong genetic effects (hazard ratios >1.5 or <0.67 in Fig. 2f).

### Mapping *Soma* loci that modulate correlations between body weight and life expectancy

Body weights were measured at 42 days (right before tails were biopsied for genotyping); and at 183, 365, 548, and 731 days (6, 12, 18, and 24 months). We used these data to map loci for body weights for both sexes so that we could align and compare body weight loci with *Vita* and *Soma* loci (e.g., Fig. 5a-c). Sample sizes for mapping *Mass* loci range from a low of 2,459 at 42 days to a high of 4,688 at 183 days (Supplementary Table 13). Models used are like those used for mapping *Vita* loci, but the variable is body weight at each of five ages.

### Actuarial analysis of the correlation between body weight to life expectancy by sex

To investigate the effects of body weight on lifespan, we adjust body weight and lifespan using the model:

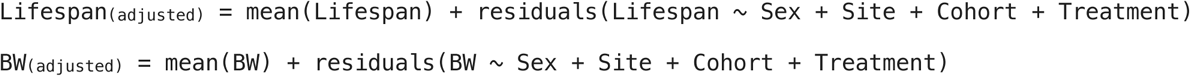

This model includes mice for which we have lifespan and weight data at one or more of the five weight ages. We computed the sex-stratified Spearman rank-order correlations between weights at 42, 183, 365, 548, and 730 days with lifespan in 15-day increments (Fig. 5d-p). We compute the –log*P* of the difference in correlation between males and females using the *cor.test* function in R. The average correlation in the combined population between adjusted body weight at 183 d (∼6 months) and adjusted lifespan is *ρ* = –0.206. When computed for males, correlation is stronger (rho = –0.284), while it is weaker for females (*ρ* = –0.110).

### *Soma* Correlated Trait Locus (CTL) mapping

We used CTL mapping^177^ to determine if a distinct set of loci modulate correlations between body weight and subsequent lifespan compared to either *Vita* or *Mass* loci. Before using this procedure, we adjusted body weight and lifespan using sex, site, cohort year, and drug treatment as covariates. At each marker we stratified survivorships based on genotypes. Ideally, the sample size for each subgroup would be above 400, but in some cases, *N* was as low as 200 due to genotype uncertainty and to the lower sample sizes at T_42_ (*N* = 2549) and T_730_ (*N* = 2208). We computed *ρ* correlations for each genotype with *N* >100 and the *z-*scores associated with differences:

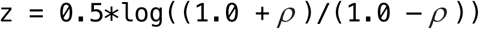

Here `cor` is a four-value vector containing the observed correlations for *CH*, *BH*, *CD*, and *BD* genotypes. We compute the sum of squares (`sumOfSq`) by multiplying the observed sample sizes (`ss`) of the *CH*, *BH*, *CD*, *BD* genotypes to the squared *z*-scores:

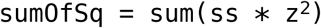

We compute the squares of sums (sqOfSum):

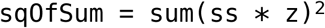

Using these values, and the sum of the sample sizes on which each correlation is based, we compute the critical value (Cv) which follows a chi-square distribution under the null hypothesis that all correlations (z-scores) are from the same distribution:

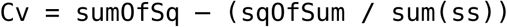

The Cv is converted to a *P* value using the chi-squared distribution using the *pchisq()* function in R, with P[X > x] and three degrees of freedom (number of genotypes – 1).

We adjust the significance threshold for multiple tests using the *p.adjust* function in R at a 5% FDR^69,178^. The 5% FDR threshold is approximately 2.75 –log*P*. While still stringent, this threshold is less harsh than that applied to *Vita* loci that use a highly conservative Bonferroni correction. Our rationale for this difference is our use of conservative non-parametric rho for comparing correlations of weight to lifespan. Differences in correlation correspond to days gained or lost relative to an average individual that can be converted to effect sizes measured in days gained or lost per gram of weight. We computed the linear regression coefficient for weight on lifespan for the four genotypes as:

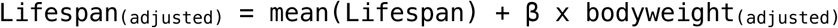

where `mean(Lifespan)` is the intercept of the total population (all genotypes combined), while the `β` coefficient is the estimated effect size of weight on lifespan based on the subpopulation defined by each genotype. This leads to effect sizes relative to an averaged mouse. There are caveats with respect to interpreting these effects because they are computed relative to this idealized mean, but values should give readers a sense of effects of *Soma* loci on life expectancies.

### Broad-sense haplotype-based heritability of *Vita* loci

We compute broad-sense genotype-based heritability by fitting a full model including the haplotype probabilities of the 29 top markers of all *Vita* loci:

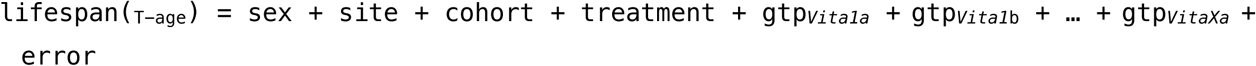

Broad-sense haplotype-based heritability is estimated by fitting this model to all survivorships. When stratifying by sex, sex is dropped from the model as well as from the H^2^_e_ component described in the computation of environmental variance component below. Computing heritability is done by taking the following approach, adapted from Falconer^179^ and Lynch and Walsh^180^ using five steps:

**Step 1**: The vector of partial total variances (PTV):

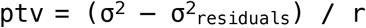

**Step 2**: An adjustment factor—the sum of the mean sum of squares for the fixed effect including the residual variance:

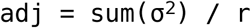

**Step 3**: The contribution of each parameter in PTV to the model:

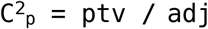

**Step 4**: The total broad-sense genotype-based heritability:

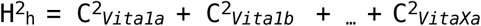

**Step 5**: The environmental variance:

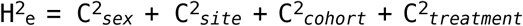

where σ^2^ is the vector containing all the means of sums of squares (including the residual) computed by the ANOVA model. σ^2^_residuals_ is the residual mean sum of squares. *r* is the average number of replicates for each genotype. H^2^_h_ can be interpreted as the broad-sense genotype-based heritability as a sum of all *Vita* loci. H^2^_e_ is the environmental variance estimate of the known environmental covariates. Unexplained variance H^2^_u_ can be computed as:

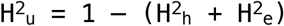

To obtain upper and lower bounds on H^2^_h_ and H^2^_e_ we generated 50 bootstrap resamples and fit the ANOVA model to each using a random subset of 90% of the survivorship. We computed median and standard deviations across all bootstraps to estimate errors of H^2^_h_ and H^2^_e_ as a function of survivorship T_age_.

### Candidate genes analysis

All annotated features located within the 95% CI were downloaded using *biomaRt*^181^. We applied several criteria to prioritize positional candidate genes: (1) first, we considered only the protein coding genes that reside in the 95% confidence intervals; (2) we then selected genes with annotated non-synonymous SNPs segregating in the population, and these were further ranked by the potential deleterious effect based on the ENSEMBL *Variant Effect Predictor* (VEP)^182^; (3) another priority score for the candidate genes was based on whether they were listed as aging and longevity genes in GenAge^183^. We also compared our positional candidate genes against known human, *C. elegans*, *D. melanogaster*, and *S. cerevisiae* genes reportedly associated with age in GenAge. To accomplish this, we use *biomaRt* to convert mouse gene symbols to the corresponding species-specific orthologous gene symbols.

Different gene sets from *Vita* loci were tested for over-representation with WebGestalt^184^ for over-representation analysis based on gene ontologies and molecular pathways (KEGG, REACTOME). We performed this analysis using three categories of gene set and background sets: 1. All protein-coding genes in *Vita* regions (*N* = 6975) versus the whole genome. 2. All protein coding genes with medium and high impact variants (as determined by the VEP) in *Vita* regions (*N* = 2502) versus all protein coding genes in *Vita* regions. 3. All protein coding genes with high impact variants (as determined by the VEP) in *Vita* regions (*N* = 409) versus all protein coding genes in *Vita* regions. Given the high number of G protein-coupled receptors (GPCRs) and olfactory receptors (OLFR) in our *Vita* regions, we repeated these three analyses while excluding GPCRs and OLFR. This reduces the size of the different gene sets to all protein-coding genes (*N* = 6112), those with either medium or high impact variants (*N* = 2354), and those with only high impact variants (*N* = 382).

### *C. elegans* candidate gene screening of *Vita9b* candidate genes

We searched for potential *C. elegans* orthologs by searching for the protein-coding genes within the *Vita9b* **locus using Ortholist2**^185^. From the list of potential orthologs, we screened C41D11.3 (ortholog of murine *Csrnp1*), F52F12.1 (*Slc22a13*), H09G03.2 (*Frmpd1*), C12C8.3 (*Trim71*), F30A10.3 (*Ip6k2*), T22C8.2 (*Hyal1*), L4440 (empty vector control) and daf-2 (positive control) from the Ahringer RNAi library^186^. The sequence-verified single RNAi clones listed above were inoculated in 10 ml LB with ampicillin (100 μg/ml) and tetracycline (50 μg/ml) overnight at 37 °C. The next day 5 ml of LB with 0.5mM isopropyl b-D-1-thiogalactopyranoside was added and inoculants were incubated for 30 min at 37 °C with shaking for pre-induction. The bacterial inoculants were centrifuged (3700 rpm for 15 min) and the pellet was resuspended in 10 ml liquid NGM with 50 µg/mL IPTG, 100 µg/ml ampicillin and 13.32 mg/ml nystatin.

Gravid adult stage TJ1060 [*spe-9(hc88)*;*rrf-3(b26)*] *C. elegans* were synchronized by bleaching. Resulting L1 stage worms were suspended in concentrated heat-inactivated OP50 bacteria in liquid NGM and transferred into U-bottom 96-well plates in a final volume of 100 µL. The worms were grown on an orbital shaker (Heidolph Titramax 1000) at 600 rpm until they reached the L4 stage (approximately 2.0–2.5 days) at 25 °C to prevent progeny production. When worms reached L4 stage, 20 µL of bacterial inoculant of each RNAi clone were added into single wells of the U-bottom 96-well plates containing worms. At least 30 worms per well and 8 wells per clone were used for screening. The MicroTracker (InVivo Biosystems) system was used to measure motility in liquid in 96-well plates by infrared beam breaks. Plates were measured daily for a period of at least 90 min.

### Exploratory data analysis, replication, and extension of UM-HET3 data by readers and students

All primary data used in this study (and more) is available at www.genenetwork.org (GN) along with tools for truncation, mapping, and basic analysis of correlations between variables such as lifespan and body weight.

## Author contributions

D.A., R.W.W., J.A., E.G.W, D.G.A., K.M.M, and R.A.M. were responsible for conceptualization. ITP colony management and informatics were directed by R.S., D.E.H., and R.A.M. Tissue and DNA sample processing and data curation was managed by S.R., D.G.A., V.D., K.L., A.C., L.L., D.A., R.A.M., and J.F.N. Statistical and bioinformatics was conducted by D.A., S.S., D.G.A., K.W.B, R.W.W, J.P.M., and M.R.M. Code development for mapping of UM-HET3 was developed by K.W.B., D.A., S.S., P.P., Z.S., and R.W.W. *C. elegan*s analysis by M.A.M, O.A.A., and S.J.M. Manuscript and illustrations by R.W.W., D.A., and E.G.W., with review and edits by all authors.

## Acknowledgments

This work was supported by grants from the NIH R01AG043930 (R.W.W.), NIH R01AG070913 (R.W.W.), NIH R01AG075813 (D.G.A.), the University of Tennessee-Oak Ridge National Laboratory Governor’s Chair in Computational Genomics for genotyping, and the University of Tennessee Center for Integrative and Translational Genomics for data processing (R.W.W., D.G.A, D.A., L.L. S.R. P.P, A.C.,Z.S.), NIH grants NIA U01AG025707 and U01AG022308 (D.E.H.), U01AG022303 and OAIC P30AG024824 (R.A.M), U01AG022307 and P30-AG13319 (R.S.), and U01AG025707 and R01GM123489 (P.P., S.S., K.W.B., R.W.W.), Ecole Polytechnique Fédérale de Lausanne (J.A.), European Research Council (AdG-787702) (J.A.), Swiss National Science Foundation (310030B160318) (J.A.), AgingX: Swiss Initiative for Systems Biology RTD 2013/153 (J.A.), NIH NIDA P30DA044223 (S.S., R.W.W.), NIH R21AG055841 and R56AG066625 (K.M.)

## Additional information

All primary data and many analytic tools associated with this papers available at www.genenetwork.org. For example, this URL will take reader directly to ITP lifespan data for 23,876 mice, including all mice analyzed in the work. https://genenetwork.org/search?species=mouse&group=HET3-ITP&type=Phenotypes&dataset=HET3-IT-PPublish&search_terms_or=10000%0D%0A&search_terms_and=&accession_id=None&FormID=searchResult Code used in mapping and analysis available at https://github.com/DannyArends/UM-HET3 Supplementary files are available upon request to: RW Williams and D Arends. Files are large collectively too large to incorporate into this bioRxiv submission. Correspondence and requests for additional material can be addressed to RW Williams.

© The Authors

---

**Extended Data 1.**
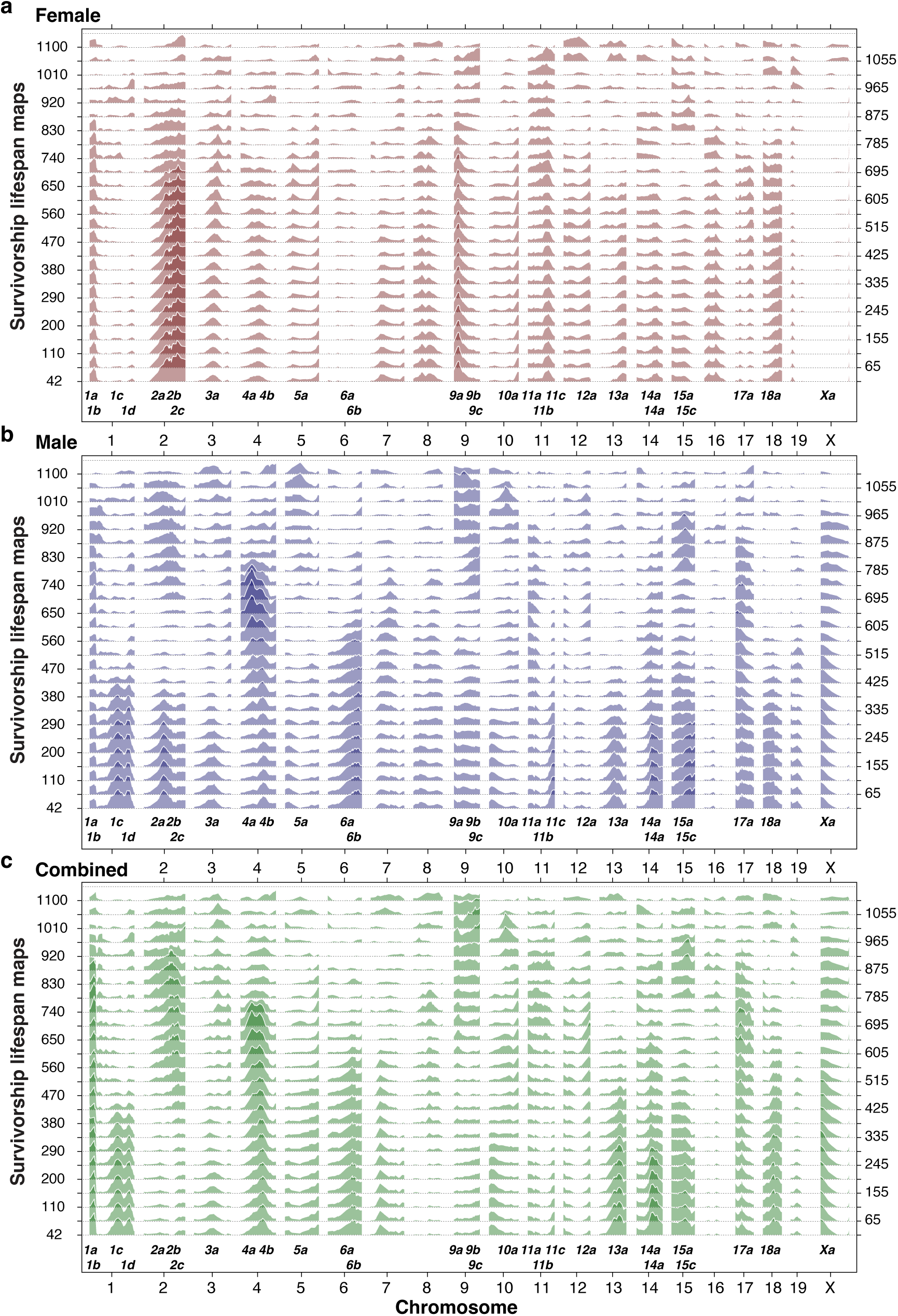
Sex-stratified and combined maps of mean lifespans of survivorships. **a,b,c,** This extends the content of Fig. 1d to included female and male survivorship maps across the entire range of survivorships. Other conventions as in Fig. 2d. The LOD threshold is defined by the horizontal dashed lines for each survivorship. Note that only every third survivorship is plotted, accounting for some *Vita* loci that do not reach significance here.

**Extended Data 2.**
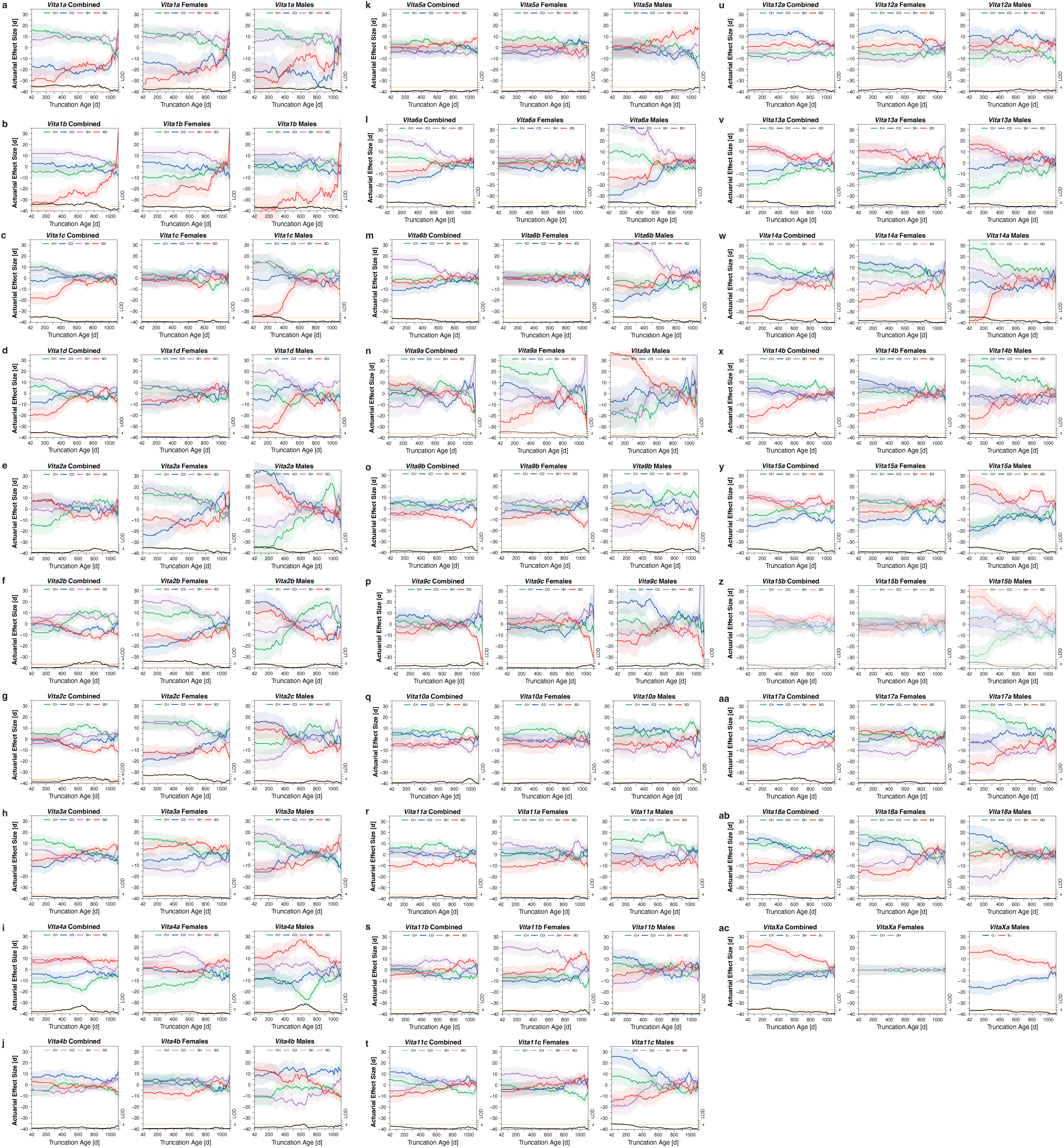
*Vita* loci genotype effect plots. Trios of effect size plots as described in detail in the legend to Figs 1 and 2. Each trio for the 29 *Vita* loci includes combined data for both sexes (left), for females only (middle), and for males only (right). To understand the genetic sources that account from timing changes in mortality rates it is helpful to compare these more complex genotype plot to pairs of haplotype plots in Extended Data 3 and 4.

**Extended Data 3.**
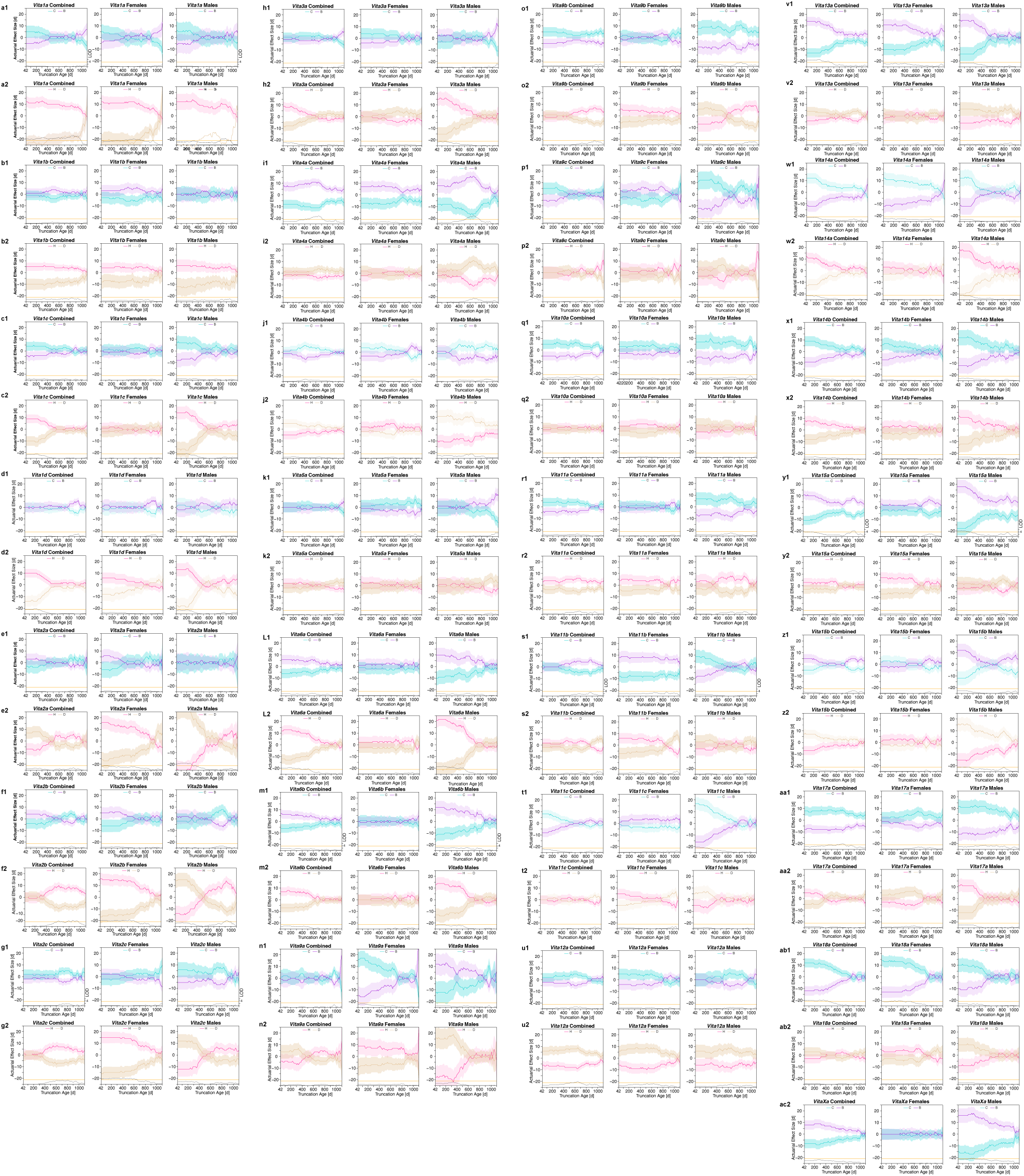
*Vita* loci haplotype effect plots. There are two trios of effect size plots for each *Vita* locus. The upper trio (blue and purple shades) give the difference in actuarial effects of the maternal haplotypes—*C* and *B*. The lower trio (pink and beige shads) give the difference in effects of the paternal haplotypes—*H* and *D*. Each These plots are symmetric and easier to interpret than the matched genotype plots in Extended Data Fig. 1. However, they generally have lower LOD scores.

**Extended Data 4.**
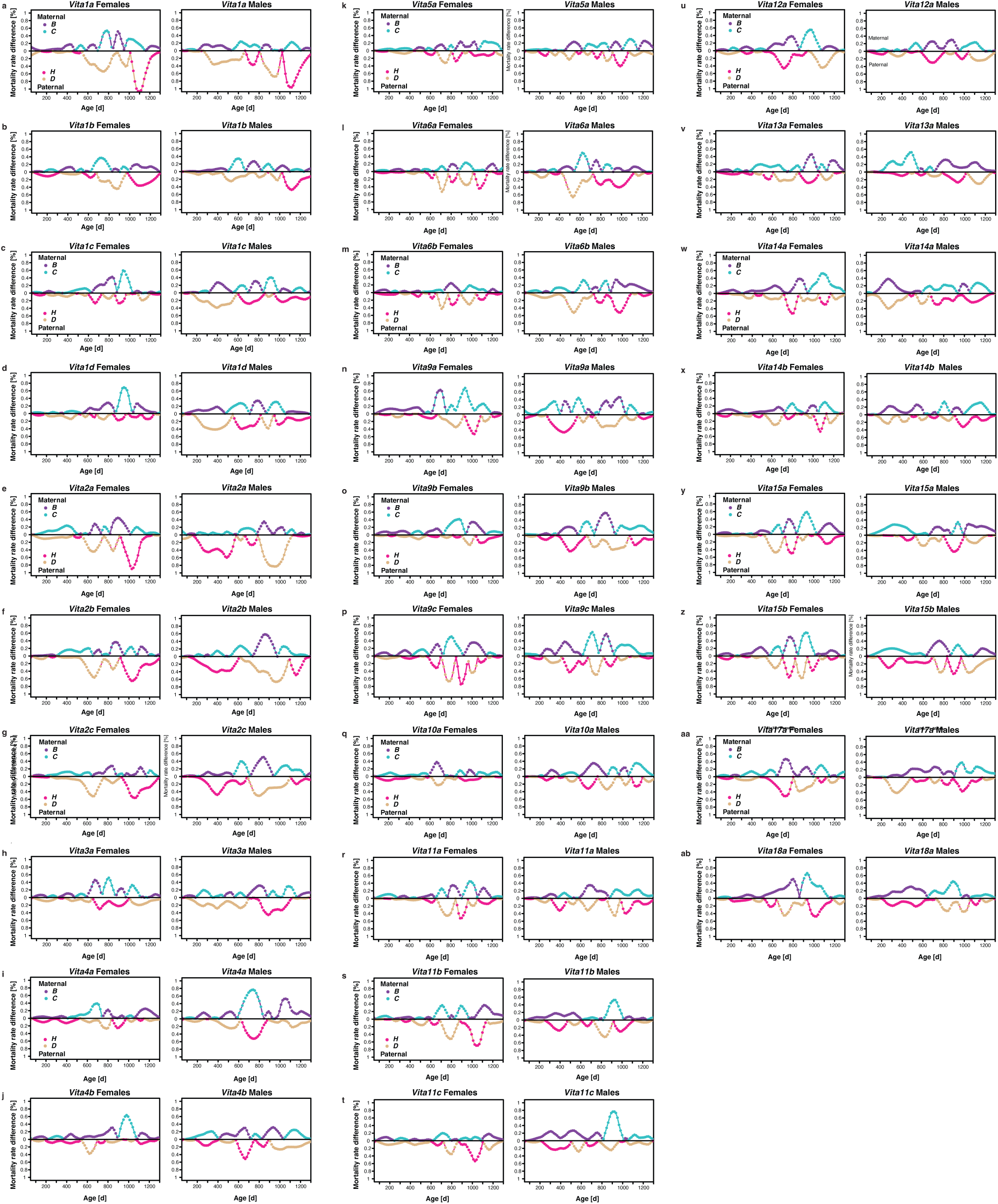
Mortality rates of maternal and paternal haplotypes of *Vita* loci. Relative mortality rate difference between the maternal pair of haplotypes (*C* and *B* on the top of each panel) and between the paternal pair of haplotype (*H* and *D* on the bottom of each panel). Here we have used a LOESS smoother with a relatively short α span 0.2. At any one age only the haplotype with the higher rate of mortality is shown, accounting for abrupt reversals.

**Extended Data 5.**
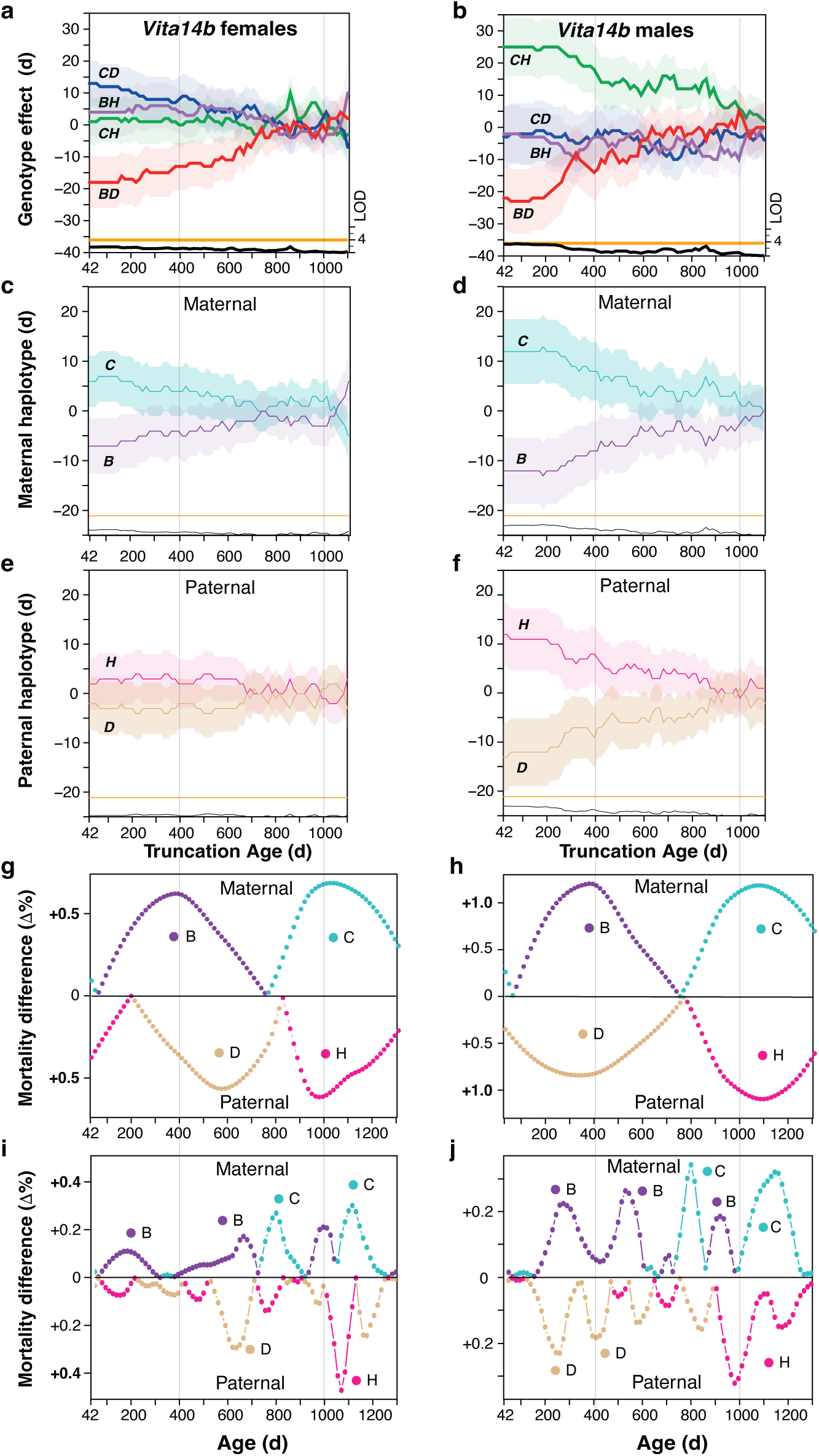
Direct comparisons of all methods of display of survivorship effect sizes of genotypes, haplotypes, and mortality rates of a *Vita14b* in females (left) and males (right). **a,b,** The actuarial genotype effect plots of *Vita14b* (see Fig. 2b) for both sexes. **c,d,** Corresponding haplotype-specific plots. **g,h,** Age-dependent relative mortality rate differences using a LOESS smoother with a span over the entire range of ages (*α* of 1) that averages mortality difference at a high level. While effects are similar between sexes, only the much stronger male effect in **b** reaches significance—the sum of the two reinforcing haplotype effects. **i**,**j**. An analysis of age-dependent differences in mortality rates using a LOESS with a span of 0.2 that exposes much finer details of both age-dependent and haplotype-dependent differences in mortality. Extended Data 4 uses only this finer-grained smoother.

**Extended Data 6.**
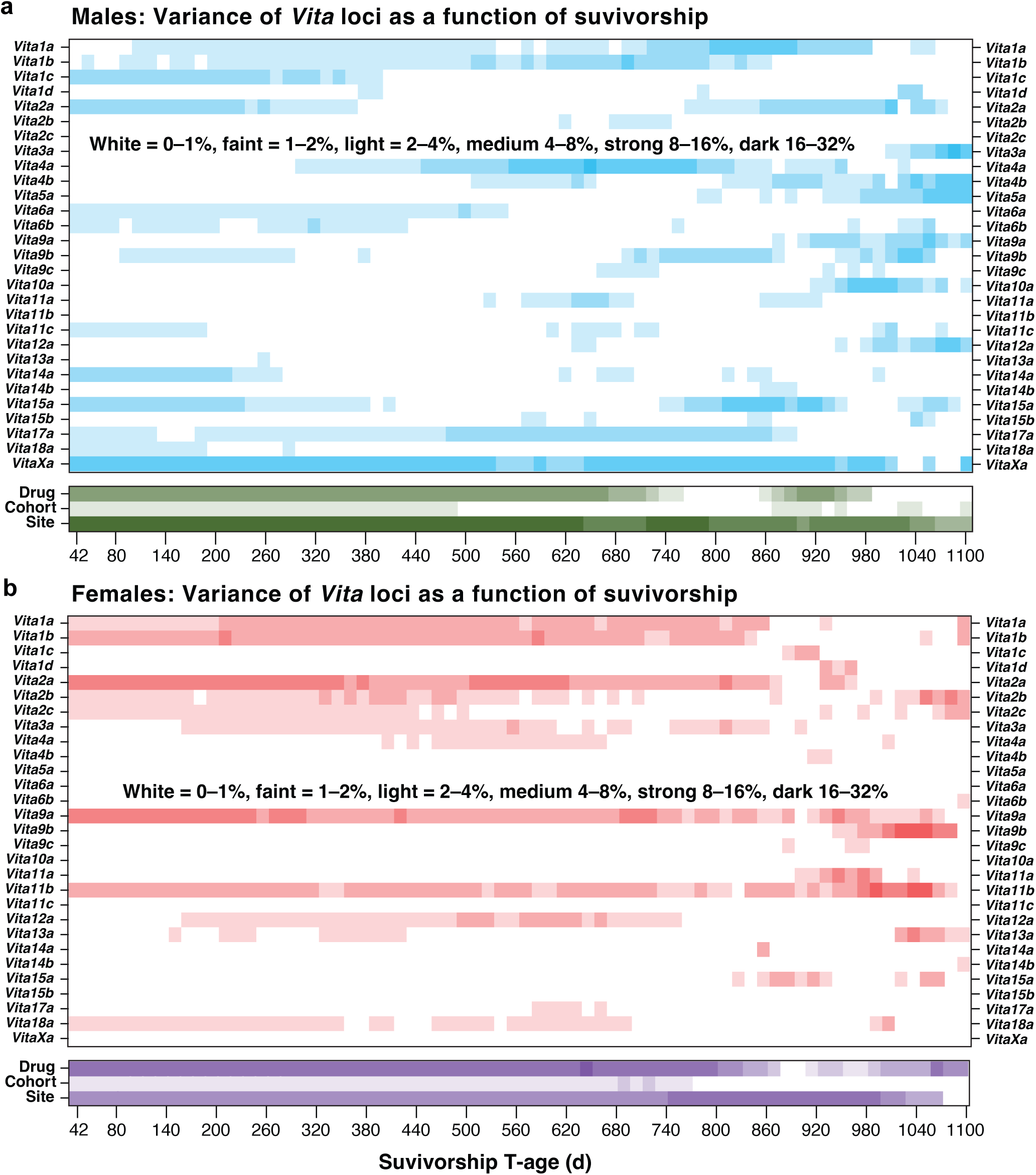
Dynamics and variance of 29 *Vita* loci as a function of sex and survivorship. **a,b,** Variance contributions of *Vita* loci were estimated for each survivorship and sex: males in **a** and females in **b** with intensity of colors indicating the approximate fraction of variance explained, where white = 0–1%, faint = 1–2%, light = 2–4%, medium 4–8%, strong 8–16%, dark 16–32%. Non-genetic experimental sources of variance (**Ve**) are provided in green for males and purple for females. **Drug** is variance attributable to multiple nominally ineffective supplementary drug treatments versus the standard chow diet. **Cohort** is variance attributable to the nine annual cycles of production of UM-HET3 mice from late spring through late fall between 2004 and 2013 (2008 was a hiatus year). **Site** is variance associated with the three ITP sites: The Jackson Laboratory, The University of Michigan, and University of Texas Health San Antonio.

**Extended Data 7.**
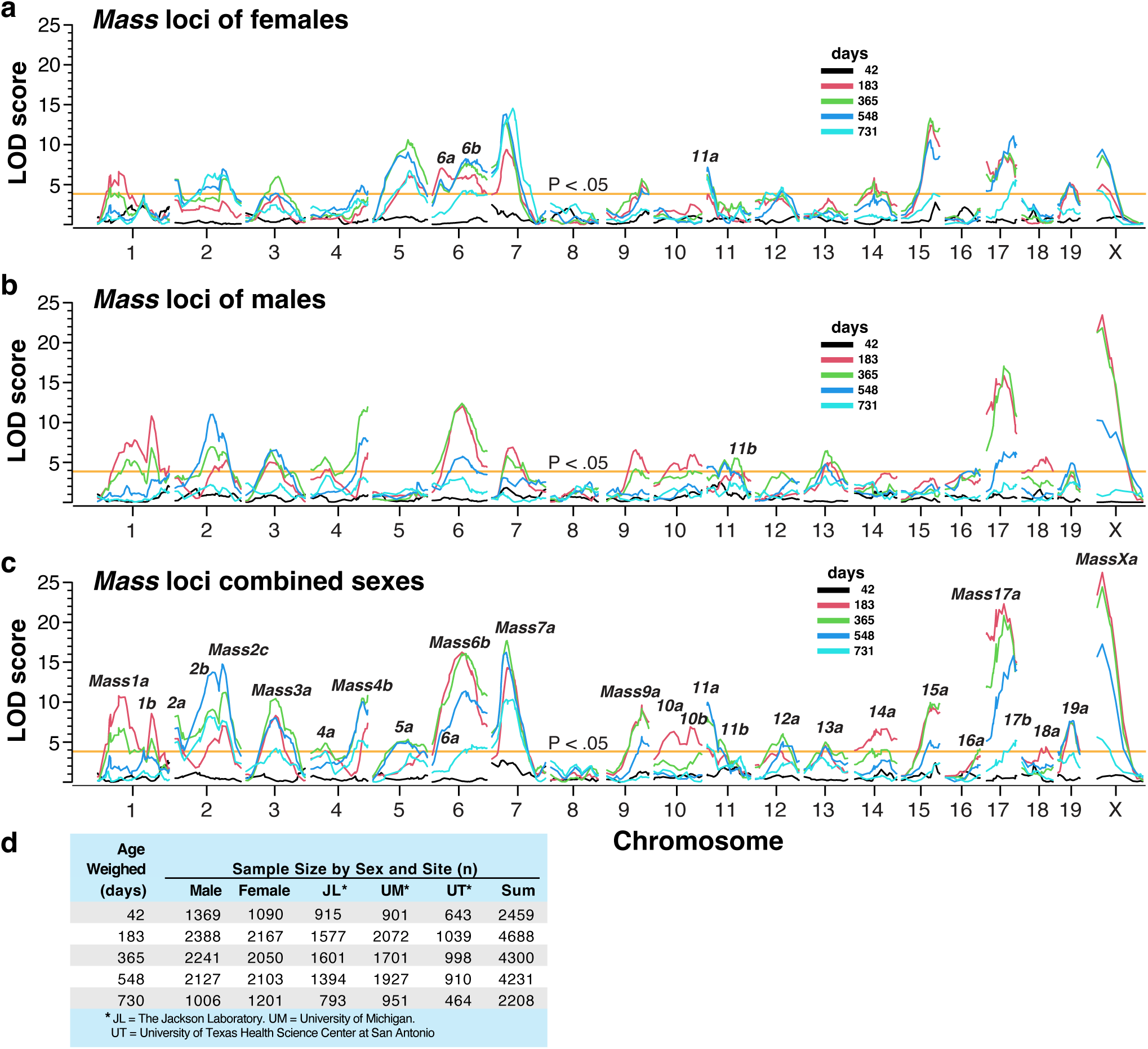
Body *Mass* loci at five ages. **a,** Female maps. **b,** Male maps. **c,** Combined maps in which all significant *Mass* loci have been labeled. Note that three of the 28 *Mass* loci are more distinct in the female map (*Mass6a*, *Mass6b*, and *Mass11a*) than in combined or male maps. *Mass11b* is more distinct in the male map. At all ages the mapping model compensates for drug treatment, cohort year, and site, and in the case of the combined sexes also adjusts for sex. Compare **(c)** with Fig. 5a, but note that color assignments are different here. **d,** Table of sample sizes for each sex and for the sites. Animals at 42 days were only weighed for the first four cohorts, accounting for the lower sample size. The lower sample size at 730 days is due in part to mortality.

**Extended Data 8.**
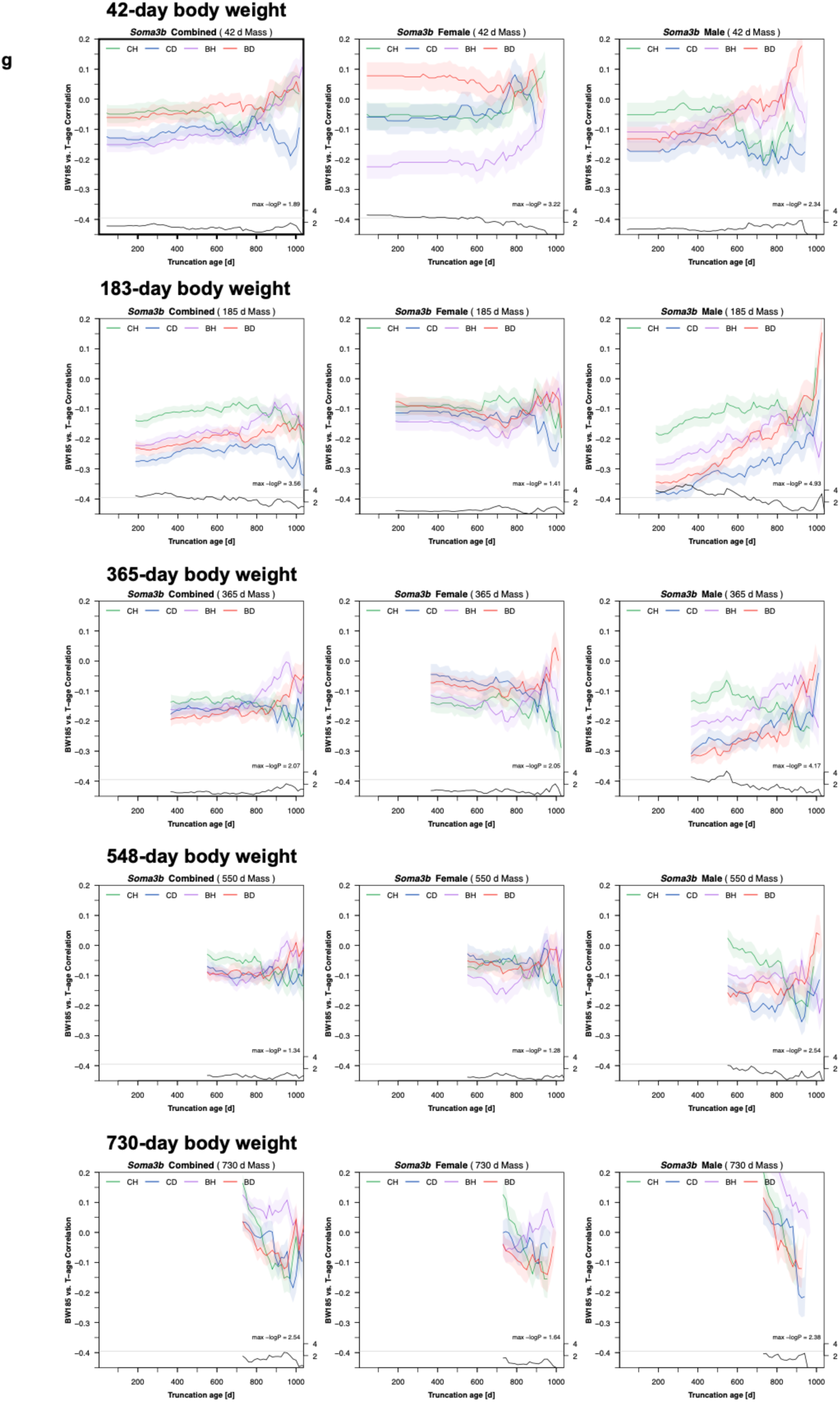
*Soma* actuarial effect size plots. **a,** An example of one of 30 *Soma* loci full actuarial effect size plots. The original figure cannot be optimized as a single-page PDF and is best viewed within Inkscape or Adobe Illustrator so that readers can examine individual or sets of *Soma* loci. The figure consists of 30 layers of loci, aligned on top of each other. Each locus is given at the five ages at which mice were weighed. In this PDF version, we have reproduced only *Soma3b* (also see Fig. 4i,j for more context). The Adobe Illustrator format file is available upon request in which we illustrate all 30 overlapping *Soma* loci.

**Extended Data 9.**
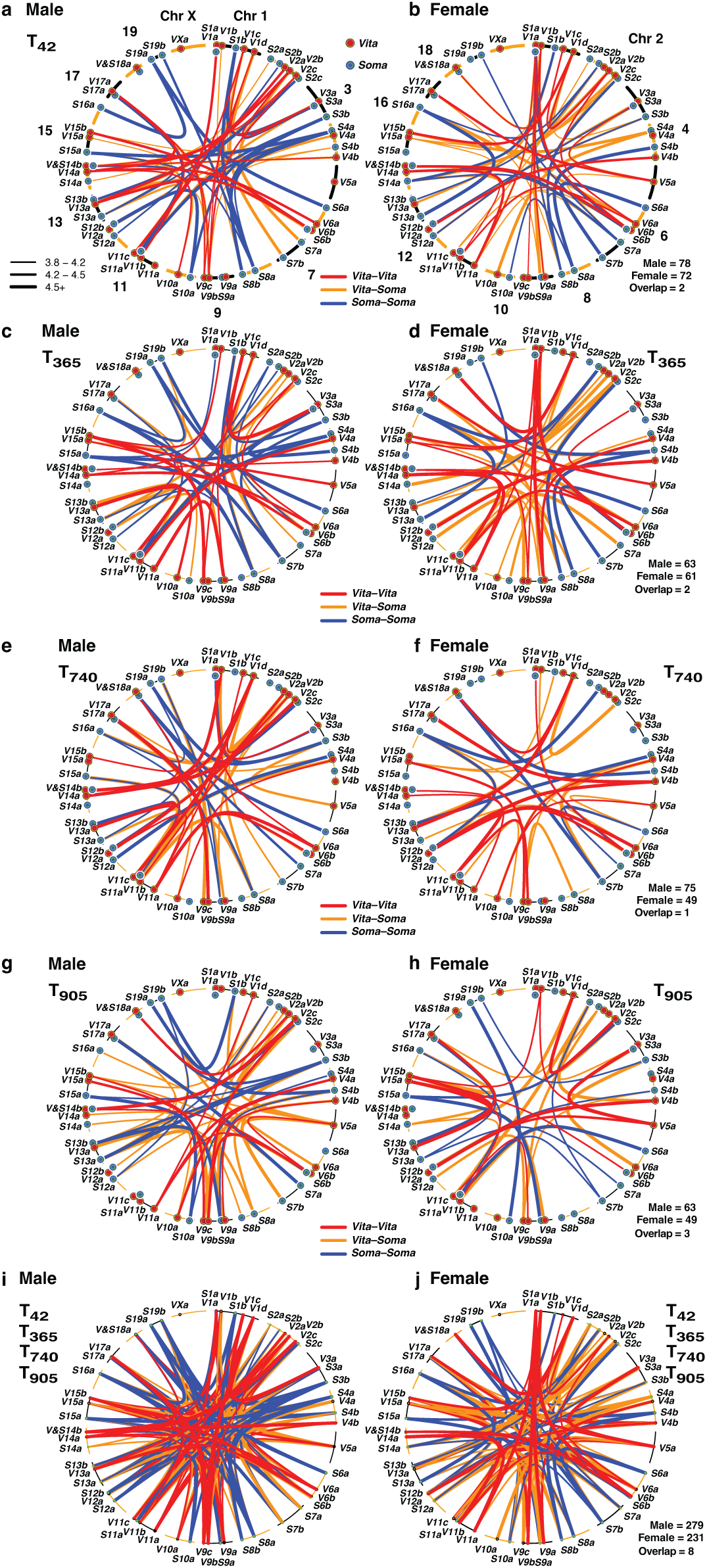
Epistatic interaction plots for the T_42_, T_365_, T_740_ and T_905_ survivorships. **a,g,** Overview of all epistatic interactions in survivorship with LODs ≥ 3.8 (thin lines), above 4.2 (medium lines), and above 4.5 (thick lines). **c–h,** Similar plots of the three older survivorships—T_365_, T_740_, and T_905_—using the same conventions. Chromosomes are labeled with abbreviated *Vita* and *Soma* symbols. Color and type of lines define partnership types (orange lines are *Vita*-*Soma* pairs). **I,j,** An overlay of all four survivorships that mainly highlights the greater cumulative numbers male than female interactions. This pair of circle plots is a set of overlapping layers useful to directly compare different survivorships and the Adobe Illustrator format is available upon request. Number of epistatic links for both males and females is given in the lower right corner of each female plot. Few epistatic interactions overlap in both sexes. All values are given in Supplementary Table 12.

